# The inflammatory micro-environment induced by targeted CNS radiotherapy is underpinned by disruption of DNA methylation

**DOI:** 10.1101/2024.03.04.581366

**Authors:** TO Millner, P Panday, Y Xiao, JR Boot, J Nicholson, Z Arpe, P Stevens, N Rahman, X Zhang, C Mein, N Kitchen, AW McEvoy, E McKintosh, G McKenna, D Paraskevopoulos, R Lewis, S Badodi, S Marino

**Affiliations:** Blizard Institute, Queen Mary University of London, UK; Barts Brain Tumour Centre, Barts Health NHS Trust, UK; National Hospital for Neurology and Neurosurgery, UCLH NHS Trust, UK; Barts Cancer Institute, Queen Mary University of London, UK; The Royal London Hospital, Barts Health NHS Trust, UK

## Abstract

Although targeted radiotherapy (RT) is integral to the increasing survival of cancer patients, it has significant side-effects, the cellular and molecular mechanisms of which are not fully understood. During RT epigenetic changes occur in neoplastic tissue, but few studies have assessed these in non-neoplastic tissue and results are highly variable. Using bulk DNA methylation and RNA sequencing as well as spatial transcriptomics (ST) in a unique cohort of patient tissue samples, we show distinct differences in DNA methylation patterns in irradiated brain tissue, whilst ST characterisation identifies specific micro-environmental niches present after irradiation and highlights neuropeptides that could be propagating neuroinflammation. We also show that in a cerebral organoid (CO) model of early changes in neurons after irradiation there are similar DNA methylation alterations and disruption of the DNA methylation machinery, suggesting that early but persistent epigenetic dysregulation plays a role in neurotoxicity. We provide a link between radiotherapy induced neuroinflammation and disruption of DNA methylation for the first time and suggest possible driving mechanisms for this chronic neuroinflammation.

## Introduction

Worldwide, there are approximately 20 million new cases of cancer per year, with almost 10 million deaths, and rates are rising[1]. Advances in primary site control has allowed increases in patient survival, and therefore the management of metastases is becoming increasingly important. The estimated incidence of brain metastasis in cancer is approximately 15%, although this is felt to be conservative[2]. Targeted radiotherapy (RT) is an essential pillar in the treatment of brain metastases (NICE guidelines), as well as vascular malformations and other focal neurological diseases, but can have significant side-effects. Short-term side effects (0-6 months) of brain irradiation can include tiredness, nausea, headaches and local hair loss, whilst long-term side-effects (>6 months) can include cognitive impairment, severe headaches (SMART syndrome), radiation necrosis, and even the development of secondary brain tumours[3]. From animal models, the mechanisms underlying RT-induced neurotoxicity have been suggested to include senescence and apoptosis of neurons, endothelial cells and glia, neuroinflammation and disruption of the blood-brain-barrier (BBB), however, the interactions of these mechanisms are yet to be fully elucidated in humans and are likely to be dynamic and complex [4, 5].

Radiation is an effective oncological treatment due to the killing of tumour cells through a potent induction of DNA damage, brought about by the direct action of radiation and the indirect action of ROS generation on DNA. Radiation is also a potent epigenotoxic stressor[6], and has been associated with changes in DNA methylation, histone dynamics, and modulation of non-coding RNAs[7]. DNA methylation alterations are the best characterised of these epigenetic changes, but reported results are highly variable, depending on the type of radiation, radiation dose, and biological context[8, 9]. The epigenetic modifications occurring in irradiated neoplastic tissue have started to be explored[10, 11], however, there are no studies to our knowledge that have assessed the changes in DNA methylation induced by ionising radiation in human brain tissue or with clinically relevant doses, and therefore the exact molecular mechanisms of radiation-induced epigenetic changes and their biological consequences for patients requires further clarification.

As the use of RT to treat CNS disease increases, it will be important to identify agents that may be able to offer some protection to normal brain tissue to preserve functional integrity and mitigate some of the challenging long-term effects of radiation damage, while not hampering the effects on tumour cells. Development of new treatment options for the protection of healthy brain within radiation treatment fields could greatly benefit from an in depth understanding of the underlying molecular mechanisms of RT-induced neurotoxicity. Here we have harnessed human brain samples from patients who have undergone targeted radiotherapy and utilised bulk DNA methylation and RNA sequencing in combination with spatial transcriptomics, as well as a cerebral organoid model system, to characterise in detail, for the first time, the disruption of DNA methylation observed in the human brain after targeted radiotherapy, the phenotypic transcriptional changes that overlie this, and to explore dysregulation in the DNA methylation machinery that are contributing to these.

## Results

To begin to explore the molecular mechanisms underlying RT-induced neurotoxicity we retrospectively identified formalin fixed paraffin embedded (FFPE) neurosurgical samples from patients that had undergone supratentorial targeted radiotherapy to a brain lesion followed by resection of the same area, and that contained peri-lesional brain tissue within 15 mm of the irradiated lesion, that would have been irradiated as part of the treatment field in our centres. Of the 14 cases identified, nine were treated for brain metastases and five for other conditions (Supplementary fig. 1A). 12 samples of peri-lesional brain tissue surrounding brain metastases from patients who had not received radiotherapy treatment were also analysed as controls. The mean time of surgical resection after irradiation was 58 months (range: 7-240 months), reflective of a range of clinically relevant timepoints.

The histological changes seen after radiation damage of the brain are well described[12] and neuropathological review found the known spectrum of changes, with reactive gliosis (Fig. 1A), chronic inflammation (Fig. 1B), vascular change in the white matter (Fig. 1B) and cortex (Fig. 1C) and areas of white matter rarefaction and necrosis (Fig. 1D) identified across irradiated brain samples.

**Figure 1:**
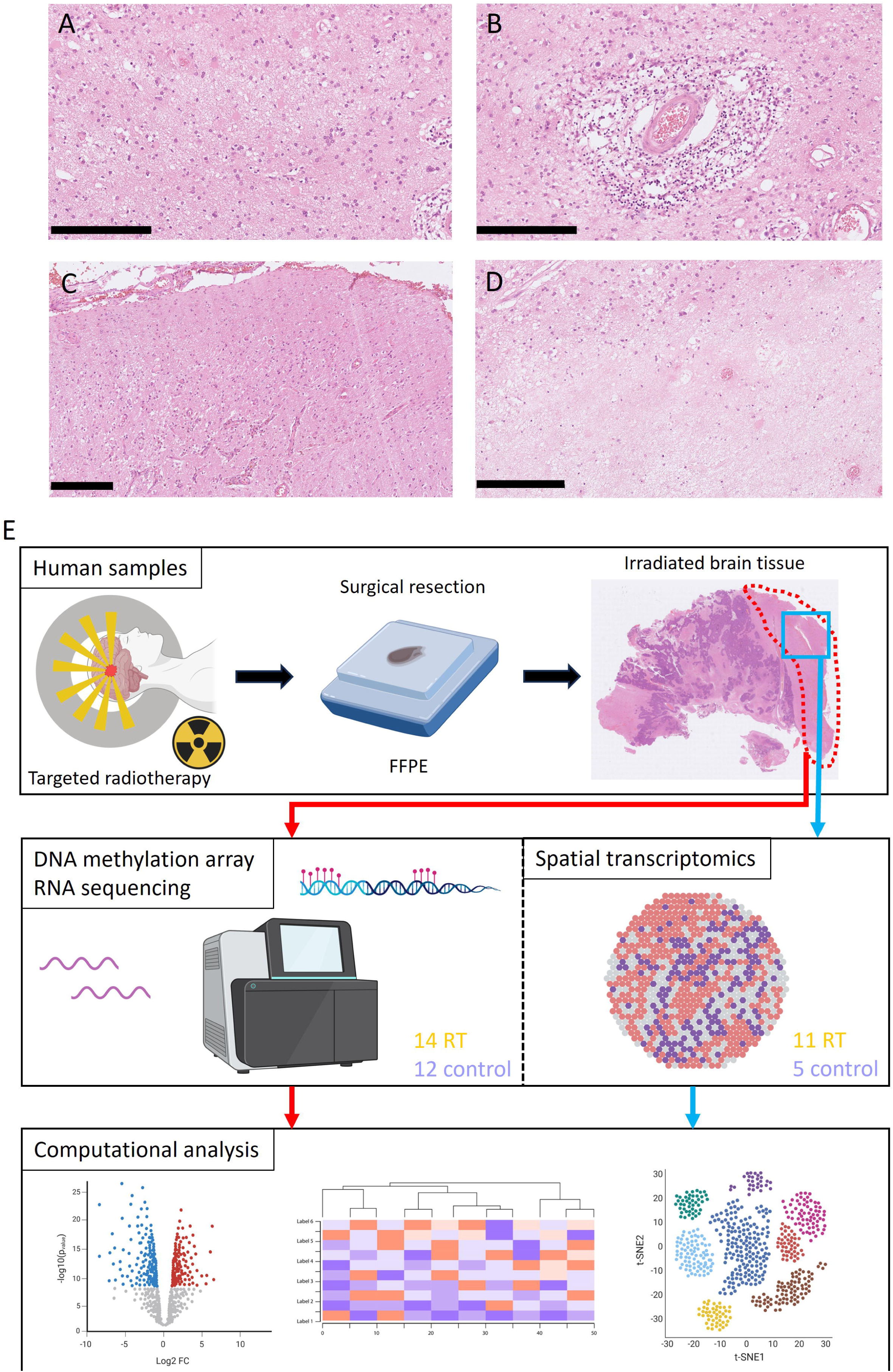
Histological characterisation and experimental pipeline. Representative histological images of irradiated brain from the study cohort are shown in **A-D**: **A** shows reactive gliosis; **B** shows white matter inflammation and vasculopathy; **C** shows cortical vessel changes; **D** shows white matter rarefaction and necrosis. Scale bars represent 250 µm. **E** Summary of the experimental pipeline for human tissue analysis; created using BioRender.

To examine the DNA methylation alterations seen in patient samples after targeted radiation, the previously irradiated peri-lesional brain tissue was micro-dissected and first characterised with DNA methylation profiling and RNA sequencing (Fig. 1E). This was supplemented with further examination of tissue from 16 cases, 11 irradiated and five controls, with spatial transcriptomics (ST; Fig 1E).

### Irradiated brain tissue harbours disrupted DNA methylation patterns

The effects of irradiation on the DNA methylation patterns of human brain tissue were studied with the use of Illumina Infinium Methylation EPIC array. Principal component analysis (PCA) of all DNA methylation array probes showed distinct clustering of irradiated and non-irradiated brain samples (Fig. 2A). Hierarchical clustering using the beta values of all differentially methylated probes (DMPs) between the groups reinforced the separate clustering observed in PCA and using these DMPs allowed annotation of a total of 4164 differentially methylated regions (DMRs; Fig. 2B). 68% (2842/4164) of the DMRs were hypomethylated in the irradiated tissue (Fig. 2C) and this tendency toward hypomethylation was observed across all genomic locations (Fig. 2D). When compared to the genomic location of probes in the array, all DMRs, both hypomethylated and hypermethylated, were enriched in promoters, exons and 5’UTR regions, whilst DMRs were depleted in 3’UTR, TEs and intergenic regions (Fig. 2E,F). These DMRs could be mapped to 4182 differentially methylated genes (DMGs).

**Figure 2:**
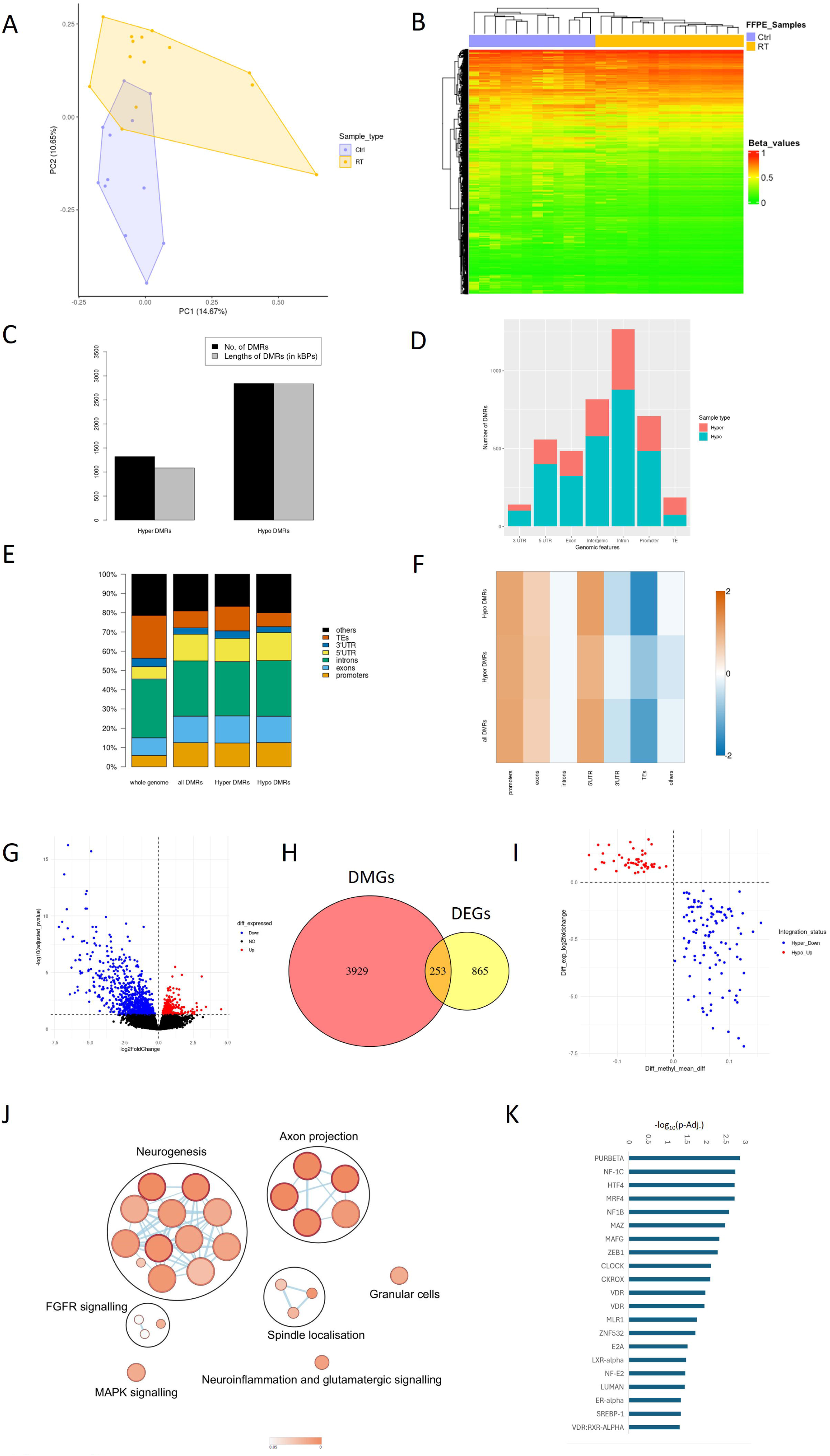
Targeted radiotherapy drives differences in the DNA methylome and transcriptome of human peri-lesional brain tissue. **A** Principal component analysis of bulk DNA methylation data using all probes after standard filtering. **B** Heatmap dendrogram for the significant DMPs identified from the FFPE samples (9468 probes in total). Both row and column clustering performed. **C** Bar plot showing number and length (in kilo-base pairs) of hyper- and hypomethylated DMRs. **D** Bar plot depicting the number of DMRs by genomic region, with proportion of hypermethylated (red) and hypomethylated (blue) DMRs. **E** Bar plot showing genomic location of DMRs as a percentage. Left hand bar is whole genome as represented on DNA methylation array. **F** Enrichment heatmap showing genomic location of DMRs (as for 2E), for all DMRs, hyper- and hypomethylated DMRs, as compared to whole genome as represented on DNA methylation array. Scale represents logFC(genomic features in DMRs / whole genome). **G** Volcano plot of the significant differentially expressed genes from RNAseq data between RT and control groups. Significant up-regulated genes are marked in red and down-regulated in blue. Non-significant are marked in black, with significance cutoff of adj. p-value <0.05. **H** Venn diagram of overlapping differentially methylated genes (DMGs) and differentially expressed genes (DEGs). **I** Scatter plot of the concordant genes identified from the integration analysis. Hypomethylated and overexpressed genes are red, hypermethylated and downregulated genes are blue. **J** GO analysis using all (53) hypo-methylated and upregulated genes from DMG and DEG overlap. **K** Table of transcription factors enriched in all (106) hyper-methylated and downregulated genes from DMG and DEG overlap using TRANSFAC database.

We then used transcriptome level data, generated from RNA sequencing from the same samples, for integration with DNA methylation data to assess the potential consequences of the DNA methylation changes on gene expression. Differential expression analysis of RNA sequencing data showed 1118 differentially expressed genes (DEGs; Fig. 2G), 78% of which had decreased expression in the irradiated brain. Integration of the two datasets showed 253 genes that overlapped between DMGs and DEGs genes (Fig. 2H). Of these overlapping genes, 159 showed concordant DNA methylation and expression (hypermethylated and low expression and vice versa; Fig. 2I). Whilst Gene Ontology (GO) analysis of all concordant genes mainly highlighted pathways involved in development, including cellular differentiation (Supplementary fig. 1B), GO analysis of genes that were hypomethylated and upregulated in the irradiated samples showed enrichment in pathways including those related to axon projection and neurogenesis, as well as FGFR signalling, MAPK signalling, and neuroinflammation and glutamatergic signalling (Fig. 2J). Chronic neuroinflammation is a critical component of radiation induced neurotoxicity[4, 5], and FGF-FGFR signalling has been suggested as an important regulator of neuroinflammation in several different cellular and pathological contexts[13], including multiple sclerosis (MS)[14], ischaemic stroke[15] and neurodegenerative diseases[16]. Specifically, FGF-1 and FGF-2 signalling has been implicated as a neuronal secreted factor that protects against neuroinflammation in animal models, likely by action on microglial migration[17, 18]. FGF-2 has also been shown to be upregulated after a single dose of radiation in a rat model[19]. The MAPK pathways act downstream of FGFR signalling and are responsible for most cellular responses to cytokines and external stress signals and are crucial for regulation of the production of inflammation mediators, including as part of neuroinflammation[20].

Interestingly, GO analysis using genes that were hypermethylated and downregulated in the irradiated samples revealed only very few broad developmental pathways, however, when these genes were searched for regulatory motifs in DNA with the TRANSFAC database[21], many potential transcription factor (TF) binding sites were revealed (Fig. 2K). Several of these transcription factors have roles in neuroinflammatory cascades. MAFG has been shown to cooperate with MAT2α in astrocytes to promote DNA methylation and repress antioxidant and anti-inflammatory transcriptional programs in MS[22]. Several studies have demonstrated the critical role of ZEB1 in promoting inflammatory responses in the CNS[23]. LXRα is important for the maintenance of the blood-brain-barrier (BBB) during neuroinflammation[24, 25], indeed disruption of the BBB is another key component of radiation induced neurotoxicity.

These findings together show that previous targeted irradiation of the brain is a strong inducer of DNA methylation alterations across the genome, with resulting DMRs showing a tendency towards hypomethylation and which are enriched in promoters, exons and 5’UTR genomic regions, suggesting a strong influence on gene expression. Integration on DNA methylation and RNA sequencing data suggest that the functional consequences of this disruption to DNA methylation appears to be related to neuroinflammation.

### Spatial transcriptomics reveals specific micro-environmental niches in the irradiated brain

Brain cell populations have a rich variation in response to any insult, and to radiation specifically, with markedly different levels of radiation sensitivity across cell-types[4, 5]. We therefore wanted to investigate the transcriptome level changes in more detail to attempt to define specific drivers of post-RT toxicity. To do this we derived spatial transcriptomics (ST) data from 16 of the cases described above (11 irradiated cases and five control cases). Haematoxylin & eosin (H&E) stained microscopy images were available for all sections and underwent neuropathological assessment for structural features. In each case an area measuring up to 6.5 x 6.5 mm^2^ was then chosen that was <15mm distant from the lesion, and therefore within the radiation field, and that contained cortex and white matter where possible (Fig. 1E; two experimental cases consisted only of white matter, one experimental case only cortical tissue, and one control case only cortical tissue). The lobar location of samples was also evenly represented across irradiated and control groups (Supplementary fig. 1A). Each region was sampled with up to 5000 spatially barcoded RNA-capturing spots with the Visium platform. For all samples, the mean number of spots under tissue was 2701, giving a total of over 40,000 individual spots, with a mean of 4436 unique molecular identifiers recovered and 17,608 genes detected per sample.

After removal of any spots containing tumour cells, by neuropathological assessment and computational filtering, 39,151 spots were included in the final analysis. Unsupervised clustering of tissue spots identified 19 clusters (Fig. 3A, B, supplementary fig. 2A). It is important to note that since ST spots here represent an area with a diameter of 55µm, and likely encompass one to several cells, each cluster represents a micro-environmental niche rather than a single cell cluster. Therefore, we next used the computational tool STdeconvolve to determine the cell-type composition of these clusters. This method can effectively recover cell-type transcriptional profiles and their proportional representation within ST spots without reliance on external single-cell transcriptomics references[26–28]. For each deconvolved cell type, annotation was performed using a combination of cluster markers and most highly expressed genes, compared with expression in known cell types using the Human Protein Atlas and STAB database[29], as well as the spatial location of cell types, confirmed by a neuropathologist (Fig 3 C, D and supplementary fig. 2A). These deconvolved cell types were further supported by correlation with the expression of known lineage markers (Fig. 3E) as well as with comparison with cell type signatures from a published dataset[30] (Supplementary fig. 2B). To further assess similarity between clusters, we computed cluster gene specificity scores for all cluster marker genes and performed Pearson correlation between cluster gene specificity signatures, which showed clusters with similar cell-type compositions were closely correlated (Fig. 3F).

**Figure 3:**
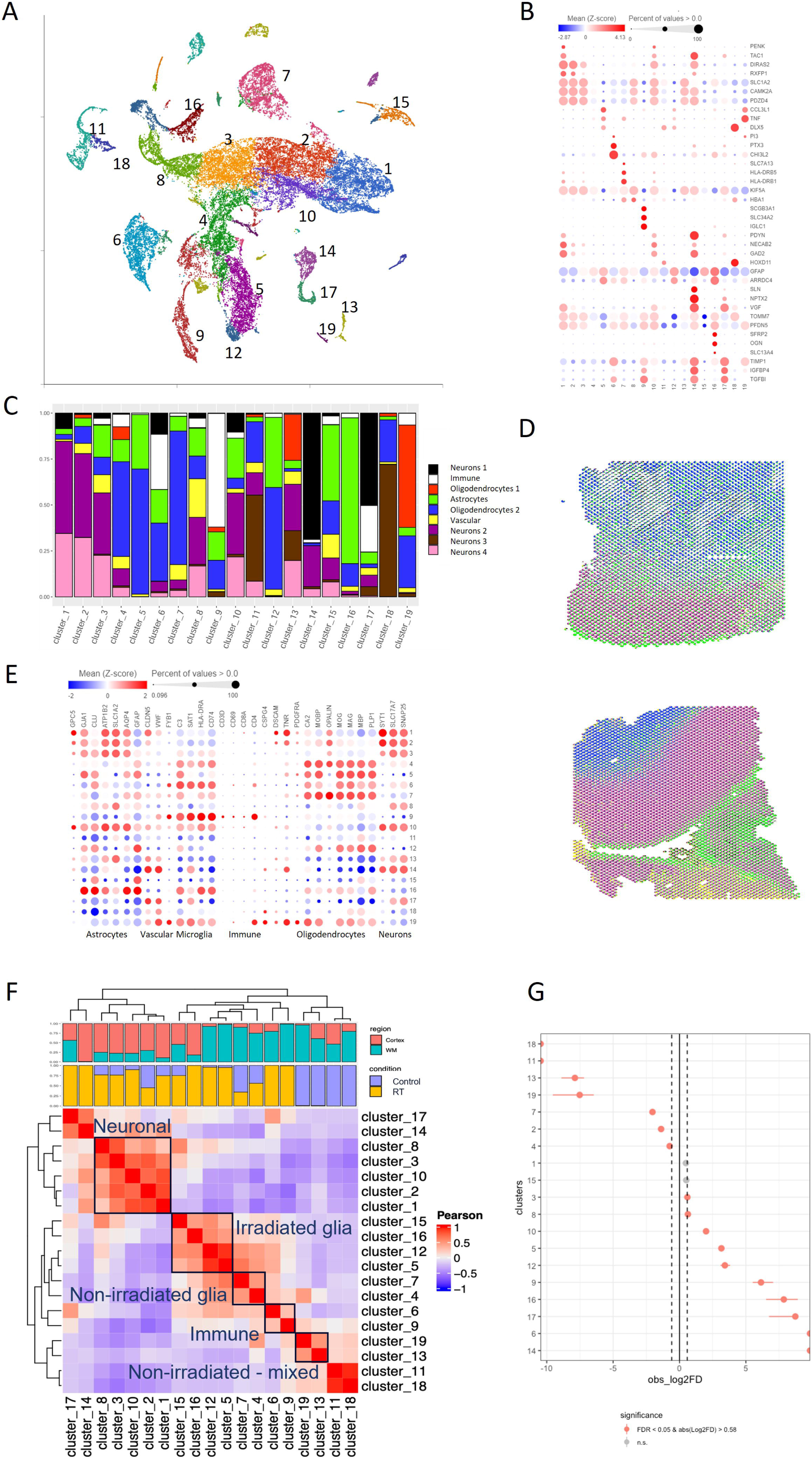
Spatial transcriptomics reveals specific micro-environmental niches after irradiation. **A** Uniform manifold approximation and projection (UMAP) plot showing all spatial transcriptomic (ST) spots after filtering (39,157 in total). Each dot corresponds to a single ST spot, coloured by cluster after unsupervised clustering. **B** Dot plot depicting selected differentially expressed genes for each cluster. Dot size corresponds to the percentage of spots expressing the gene in each cluster, and the colour represents the average expression level. **C** Bar pot depicting proportion of deconvoluted cell types in each cluster using STdeconvolve. **D** Two representative spatial plots showing deconvolved cell type distribution across sample. Each pie chart represents a ST spot and proportion of the pie chart represents deconvolved cell type proportions (see also supplementary fig. 3A). Colours are as for 3C. **E** Dot plot depicting selected lineage marker genes for each cluster and associated cluster labelling. Dot size corresponds to the percentage of spots expressing the gene in each cluster, and the colour represents the average expression level. **F** [30] Clustered correlation plot for cluster marker genes and bar plots of proportion of cortex / white matter location and experimental group for spots within each cluster. Groups of clusters are annotated as per main text. **G** Dot plot showing results of proportion test, to test for significant spot enrichment of cluster for experimental group. Enrichment in RT to the right and enrichment in control to the left.

These combined data were then used to annotate micro-environmental niches. Clusters 1, 2, 3, 8, 10 were made up predominantly of neurons and were labelled as neuronal micro-environmental niches. Within these, cluster 2 was highly enriched in control spots and cluster 10 was highly enriched in irradiated spots (Fig. 3F, G and Supplementary fig. 2C). Within these neuronal niches there were also smaller number of astrocytes and oligodendrocytes. Clusters 4, 5, 7, 12, 15, 16 were predominantly composed of oligodendrocytes and astrocytes. Within this glial grouping, clusters 4 and 7 were enriched for control spots, whilst 5, 12 and 16 were enriched in irradiated spots (Fig. 3F, G, supplementary fig. 2C). These clusters were therefore labelled as irradiated and control glial niches. Clusters 6 and 9 had a prominent immune cell component, and therefore were assigned as an immune active niche. Cluster 13 and 19 displayed mixed cell types enriched in control spots and both of these clusters were each primarily composed of spots from one sample (Cnt11 and Cnt3, respectively). Clusters 14 and 17 were almost entirely from one single sample (RT4), as were 11 and 18 (Cnt2). Therefore, these clusters have not been included in our downstream analysis as any findings from these clusters would be difficult to generalise.

Clustering and annotation of micro-environmental niches has allowed us to identify those specific to the irradiated brain and provided neuronal and glial niches to compare between irradiated and non-irradiated brain, as well as niches only seen in irradiated brain composed predominantly of inflammatory cells.

### Irradiated glial niches have an inflammatory and angiogenic phenotype

To characterise the biology of specific clusters we next performed GO analysis using the cluster markers for each cluster of interest (supplemental data). Clusters that were not significantly enriched in either irradiated or control spots, or that were predominantly composed of spots from a single sample were not examined further (Fig. 2G, supplementary fig. 2C, D). We first explored the non-irradiated glial clusters 4 and 7, in which oligodendrocytes were the most highly represented deconvolved cell type, with a smaller contribution from all other cell types (Fig. 3C). GO analysis using the cluster markers of both of these clusters highlighted pathways that were involved in gliogenesis and oligodendrocyte function (Supplementary fig. 3B, C). This analysis also showed pathways involved in immune responses and inflammation were prominent in cluster 4, whilst a number of developmental related pathways, including those linked to neurogenesis were associated with cluster 7. Cluster 12 was highly enriched in spots from irradiated cases and was composed almost entirely of oligodendrocytes and astrocytes by deconvolution, with oligodendrocytes predominant (Fig. 3C). GO analysis using markers of cluster 12 highlighted similar pathways to the non-irradiated glial niches, with pathways involved in nervous system development, gliogenesis and myelination all featuring, as well as inflammatory pathways, leucocyte migration and cytokine activity (Supplementary fig. 3D).

Neuroinflammation involving glia is known to be a feature of the micro-environment in the setting of brain metastases[31, 32], explaining the presence of inflammatory pathways in the non-irradiated glial niches, but to decipher which mechanisms were preferentially activated after targeted radiotherapy, we next examined genes that were overexpressed specifically in the irradiated glial niche after calculating DEGs between clusters (Fig. 4A, C). Pathways that were enriched in cluster 12 in this comparison included those related to signalling and development, but also those involved in inflammation and interestingly, angiogenesis (Fig. 4B, D). Examples of genes that were significantly overexpressed in the irradiated glial niche from the comparisons with both non-irradiated glial niches and involved in the pathways described above included *AGT*, as well as FOS family genes and *GADD45*, neither of which to our knowledge have been previously directly associated with radiotherapy. *FOS* is an immediate early gene that can act as a sensor of inflammation in the brain. It has been most well studied in association with neuronal activity, however, there is evidence in animal models that it is also expressed in astrocytes, oligodendrocytes and microglia[33]. GADD45 is a demethylase involved in DNA damage repair, participating in BER of mutated cytosines, possibly through interaction with DNMT1 and DNMT3A. It is widely expressed in the nervous system where it is required for the stress response, apoptosis, and mitosis of neuronal and glial cells[34]. BBB disruption is a major component of the brain’s response to radiation. AGT is the main precursor of all angiotensin peptides, which are involved in vascular remodelling. Angiotensin converting enzyme (ACE) inhibitors are effective countermeasures to chronic radiation injuries in rodent models and there is some evidence for similar effects in humans from retrospective studies of injuries in radiotherapy patients[35].

**Figure 4:**
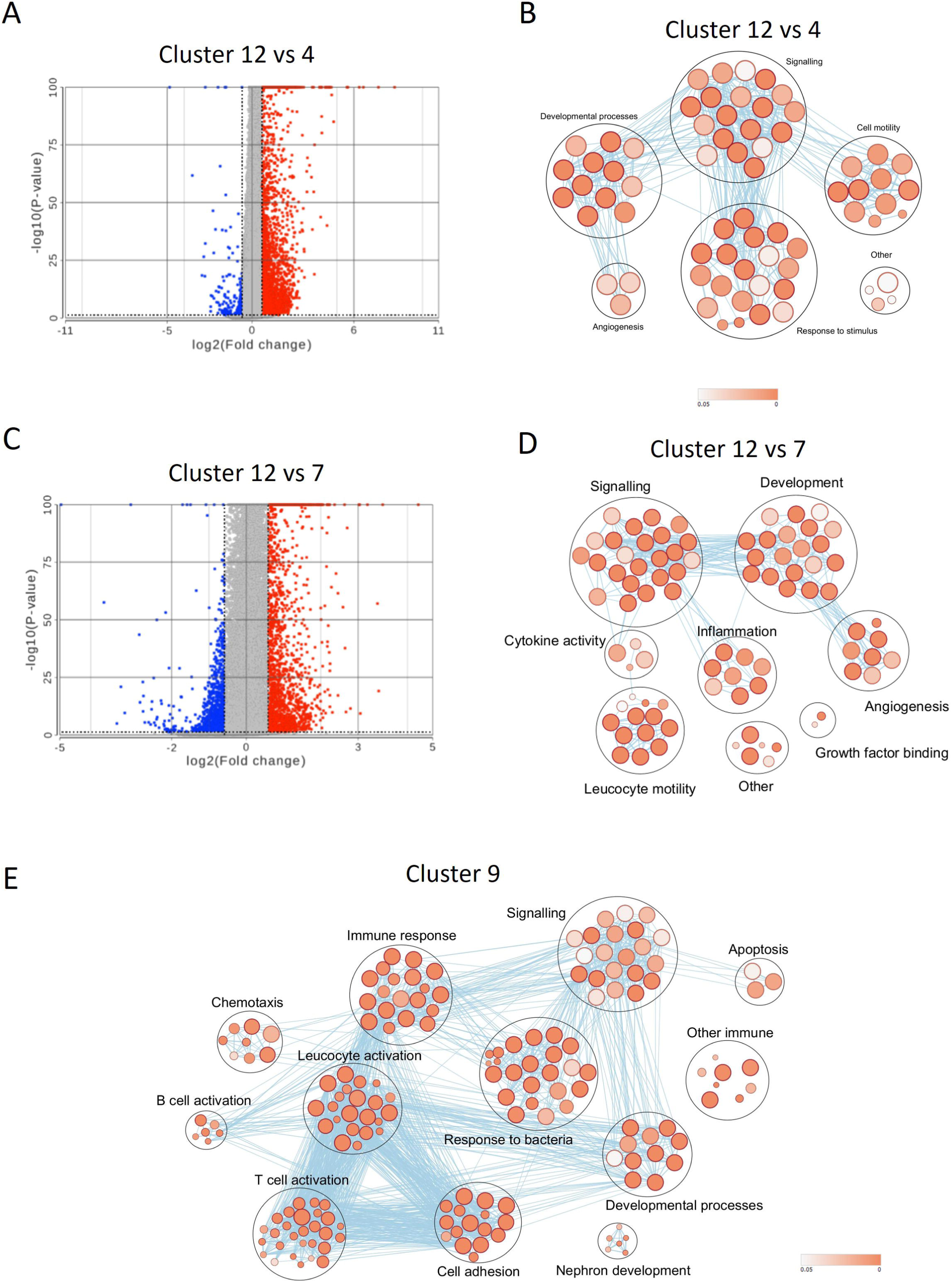
Exploration of irradiated and non-irradiated glial niches shows an inflammatory and angiogenic phenotype after irradiation. **A** Volcano plot showing differentially expressed genes between cluster 12 (irradiated glial) and cluster 4 (non-irradiated glial). Blue dots are significantly overexpressed in cluster 12 and red are significantly overexpressed in cluster 4. Significance threshold is -1.5<FC>1.5 and p-value <0.05. **B** GO analysis using 150 genes most highly expressed in cluster 12 when compared to cluster 4. **C** Volcano plot showing differentially expressed genes between cluster 12 (irradiated glial) and cluster 7 (non-irradiated glial). Blue dots are significantly overexpressed in cluster 12 and red are significantly overexpressed in cluster 7. Significance threshold is -1.5<FC>1.5 and p-value <0.05. **D** GO analysis using 150 genes most highly expressed in cluster 12 when compared to cluster 7. **E** GO analysis using top 150 cluster marker genes for cluster 9.

The deconvolved cell composition of cluster 9 had a significant proportion of immune cells (Fig. 3C) as well as a contribution from astrocytes and oligodendrocytes. Given the predominance of these immune cells it was felt there was no cluster for direct comparison in the non-irradiated brain for this cluster. It is likely that this reflects that the chronic inflammatory cell infiltrate seen after irradiation is much more prominent than in non-irradiated brain harbouring metastases and it may be that the glial cells that are part of this cluster are those most involved in immune cell infiltration. GO analysis using cluster makers for this immune active cluster highlighted pathways that were heavily enriched for those involved in inflammatory processes, reflecting a broad inflammatory response, and T and B lymphocytes specifically (Fig. 4E). The chronic inflammation brought on by irradiation is thought to be mediated through microglia to then involve macrophages and lymphocytes[4, 36], and whilst our exploration of this immune active cluster would be consistent with this, we are not able to resolve these different immune cell types individually at present.

Characterisation of irradiated glial niches with spatial transcriptomics supports roles for chronic inflammation and disruption of brain microvasculature after irradiation, involving both immune cells and glia.

### Irradiated neurons secrete a pro-inflammatory cocktail of neuropeptides

Neuroinflammation is essential for the resolution of CNS injury after exposure to many insults, including infection, trauma, ischaemia and others. The severity of the neuroinflammatory response, and its functional consequences, are proportional to the type and extent of damage. This response can have both beneficial (axonal repair, clearance of infection) and detrimental effects (injury to neurons, demyelination). This damage eventually leads to the clinical symptoms experienced by patients after radiotherapy. The prevailing theory is that neuroinflammation after radiotherapy is driven in a large part by the response to neuronal damage, which then contributes to activation and senescence of astrocytes, microglia and endothelial cells[4, 5], which themselves have also been affected by radiation damage. These damaged neurons secrete pro-inflammatory factors, which are currently not fully identified, however, elucidation of these will be essential for any efforts to protect healthy brain from the neuroinflammatory sequelae post-radiotherapy. Therefore, we next focused in more detail on irradiated neuronal niches.

Of the niches identified with a prominent neuronal component (Supplementary fig. 4A), cluster 2 was highly enriched in control spots and cluster 10 was highly enriched in irradiated spots (Fig. 3G). GO analysis using cluster markers showed that neuronal pathways were prominent for both clusters (Supplementary fig. 4B, C), further supporting our annotation. From our reference-free deconvolution, four separate neuronal subtypes were present, however, the distribution of these was relatively uniform across neuronal clusters (Fig. 3C), whilst cluster 10 had a larger proportion of astrocytes, possibly reflecting a reactive astrocyte population. The contribution of spots from different brain lobes to the neuronal niches was also consistent across niches (Supplementary fig. 4D).

As an initial comparison between the irradiated and non-irradiated neuronal clusters, all spots within these clusters were scored against signatures from published datasets for broad biological responses. The irradiated cluster 10 score more highly for signatures for senescence[37] and inflammation[38]. Conversely, non-irradiated cluster 2 scored more highly for a signature for intact neurons that was found to be inversely associated with likelihood of neurodegeneration in multiple sclerosis[39], a disease that shares several features with radiation-induced brain injury (including chronic inflammation, increased ROS, white matter pathology and long-term cognitive decline; Fig. 5A). We then assessed clusters against specific elements of the inflammatory cascade (immune checkpoints[40], cytokine response[41], HLA association[42] and inflammasome activation[41]) and found that cluster 10 scored more highly for each signature (Fig. 5B).

**Figure 5:**
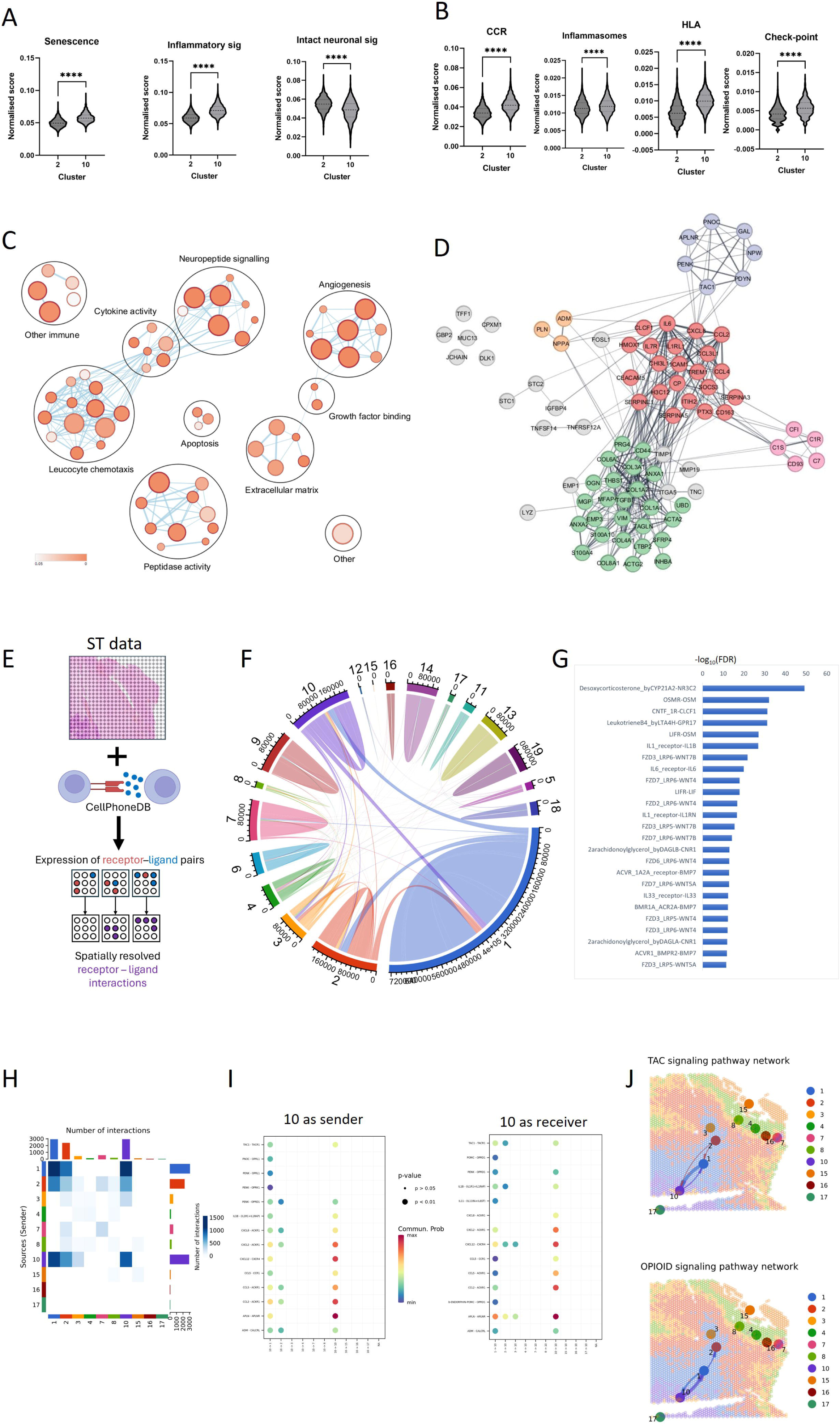
Irradiated neurons are residing in an inflammatory micro-environment with active neuropeptide and cytokine signalling. **A** Violin plots for signature scores of all spots in cluster 2 (non-irradiated) and cluster 10 (irradiated) for senescence, inflammation and intact neurons (see main text for details). Y axes show z-score normalised against gene expression per spot. Significance is tested with unpaired t-test. **B** Violin plots for signature scores of all spots in cluster 2 (non-irradiated) and cluster 10 (irradiated) for components of the inflammatory pathway (see main text for details). Y axes show z-score normalised against gene expression per spot. Significance tested with unpaired t-test. **C** GO analysis using 100 genes most highly expressed in cluster 10 when compared to cluster 2. **D** String network of 100 genes most highly expressed in cluster 10 when compared to cluster 2. Colours represent clusters from clustering performed in Cytoscape String app, with clustering performed with a granularity value of 4. **E** Schematic of generation of receptor-ligand (R-L) interaction scores. ST data was interrogated using a curated R-L database (CellPhoneDB [44]) to generate interaction scores where receptors and ligand were expressed in close proximity (also see Methods). **F** Circos plot showing interactions between all clusters using method shown in 5E. Chord thickness represents number of R-L interactions. **G** Bar plot showing 25 most significantly different R-L pairs when interaction scores were compared between cluster 2 and cluster 10. Interaction scores between clusters were compared using a Wilcoxon Test and p-values adjusted for multiple hypothesis testing using the Benjamini-Hochberg procedure. **H** Heatmap created using CellChatV2 showing number of R-L interactions between each cluster in Cnt8. **I** Bubble plots created using CellChatV2 showing R-L communication probability for selected R-L pathways in Cnt8. Left panel is for cluster 10 is the sender (ligand is expressed by cluster 10); right panel is for cluster 10 as receiver (receptor is in cluster 10). **J** Spatial plots for TAC signalling and opioid signalling created using CellChatV2 showing R-L interactions between each cluster in Cnt8. Thickness of arrows represents number of interactions.

We then identified genes that were differentially expressed between these irradiated and non-irradiated clusters (Supplementary fig. 4E). GO analysis using these DEGs revealed pathways enriched in the irradiated cluster that included neuropeptide signalling, cytokine activity and leucocyte chemotaxis as well as angiogenesis, growth factor binding and extracellular matrix related pathways (Fig. 5C). Visualisation at an individual gene level of the genes most highly expressed in the irradiated neuronal cluster 10 showed a close connection between several neuropeptides (*TAC1*, *PNOC*, *PENK*, *APLNR*, *PDYN*, *GAL*, *NPW*), and of these, *TAC1*, *PENK*, *PDYN* and *PNOC* had associations with components of the inflammatory cascade that is known to drive neuroinflammation (*CXCL8*, *IL6* and *CCL2*; Fig. 5D). Tachykinins and opioid peptides, the neuropeptides produced by *TAC1* and *PENK*, are known to be involved in communication with the immune system, although neither have been previously implicated in the radiation response specifically. *TAC1* was also associated with *ADM* and *NPPA*, genes with known roles in blood vessel homeostasis.

When we integrated our DNA methylation data from bulk sample analysis with the 588 genes that were found to be differentially expressed in the comparison of irradiated vs non-irradiated neuronal clusters, we found that *TAC1* and *PENK* were among the 74 concordant genes, suggesting that DNA methylation may be involved in the regulation of these genes in the post-irradiation brain.

ST data can be interrogated using receptor-ligand (R-L) interaction databases to examine the interactions controlling disease mechanisms directly within tissue[39, 43]. Therefore, to associate validated disease pathways with intercellular communication, we sought to identify R-L interactions that were altered between the irradiated and non-irradiated neuronal clusters. We first analysed the expression of nearly 1500 curated R-L pairs from CellPhoneDB[44, 45] in our ST data, deriving interaction scores based on the mean expression values of R-L pairs expressed in close proximity to each other (Fig. 5E and Methods). This analysis generated spatially resolved expression data for 579 R-L interactions and allowed the assessment of communication between clusters across all samples. Interactions within clusters predominated over interactions between clusters when considered across all samples (Fig. 5F). It should be noted here that since spots of the same cluster are usually adjacent, the potential for interactions within clusters outweighs that for interactions between clusters, thus, communication within the milieu of each niche will likely reveal important interaction patterns. However, there were substantial interactions between the group of neuronal clusters (1, 2, 3, 8 and 10), as well as smaller numbers of interactions between most clusters. We next used this method to test the differences between the communication patterns of cluster 2 (non-irradiated) and cluster 10 (irradiated). 125 R-L interactions had significantly different expression scores between these two clusters, and within these 125 R-L pairs, roles in inflammation and WNT pathway signalling were prominent (Fig. 5G). WNT signalling has roles in many processes including neuroinflammation[46, 47]. This is intriguing as even though cluster 10 is mainly interacting with other neuronal niches, inflammatory signalling is still prominent, suggesting that this signalling could be driving inflammation in other neuronal niches.

We next harnessed another method used for integrating ST and R-L data, CellChat v2[48–50], to examine some of the pathways identified from our comparisons above. CellChat v2 uses a simplified mass action-based model, which includes the core interaction between ligands and receptors with multi-subunit structure along with modulation by cofactors and integrates this with spatial location data. It uses a database of ∼3300 interactions incorporating CellPhoneDB and other annotated databases, including neuron specific R-L interactions (NeuronChatDB[51]). In samples that made significant contributions to cluster 10 (Supplementary fig. 2D), we assessed interactions between clusters and analysed specific neuropeptide signalling and directly connected inflammatory pathways to check whether these pathways were involved in active signalling by cluster 10. Within these samples we again saw that cluster 10 was mainly interacting with neuronal niches (Fig. 5H, supplementary fig. 5), but also with glial (Supplementary fig. 5A, D) and immune (Supplementary fig. 5A, B) niches. Cluster 10 appeared to be communicating using TAC and opioid peptide R-L interactions (Fig. 5I, J and supplementary fig. 5A-D) as well as known inflammatory interactions (Fig. 5I and supplementary fig. 5A-D).

This direct comparison between irradiated and non-irradiated neuronal clusters has shown that irradiated neurons are residing in a highly inflammatory micro-environment that is likely being actively maintained by a combination of neuropeptide and cytokine signalling.

### Cerebral organoid model shows DNA methylation disruption, neuropeptide upregulation and dysregulation of methylation machinery

The human tissue that we have examined has been resected with a latency of at least several months after RT, and we next wanted to evaluate methylation changes that might be occurring immediately after RT as well as the potential consequences and causes of these changes. Some of the disruption of methylation in patient samples will be due to downstream cascades brought on by the brain’s long-term inflammatory response to irradiation[52, 53] but there is likely an important contribution from changes to DNA methylation that take place at the early timepoints after irradiation and therefore could be amenable to drugs that could act as an early barrier to the development of neurotoxic phenotypes. To examine changes in DNA methylation and their underlying mechanisms at early time points after irradiation, we used a cerebral organoid (CO) model, a system which allows human-specific modelling of complex structures and functions of the brain[54].

For this model, COs were derived from patient-derived EPSCs[55] and irradiated at day 48 of development with a radiation dose of 24 Gy. Since we are interested in the early neuronal response, we chose this timepoint as at this time, within the neuropil areas between rosettes, our COs contain neurons that display markers of maturity (NeuN, TBR; supplementary fig. 6B-D) in a complex environment but have a limited non-neuronal population[56, 57]. 24 Gy is a dose matched to that which patients receive as part of routine targeted radiotherapy. To assess DNA methylation differences, we irradiated COs and then performed DNA methylation profiling after four weeks.

As for the patient samples, PCA of all DNA methylation probes showed distinct clustering between the two groups (Fig. 6A). Annotation of DMRs showed that 54% were hypermethylated, this distribution between hypo- and hypermethylated DMRs was more evenly shared than in the patient samples (Fig. 6B), which had shown a strong hypomethylation bias. As with the patient samples, the ratio of hypo-to hypermethylated DMRs was broadly reflective of the overall distribution of DMRs within promoters, exons and introns, TEs and other regions, whereas 5’UTR regions were more hypermethylated (66% hypermethylated DMRs) and 3’UTR regions were more hypomethylated (31% hypermethylated DMRs) (Fig. 6C) than the overall proportion. The DMRs, both hypo- and hypermethylated were also preferentially found in promoters, exon and 5’ UTR regions (Fig. 6D), as was the case in patient samples. We then mapped the CO DMRs to DMGs and when we compared DMGs from patient samples to those from our CO model, there were a considerable number of genes shared (844) between the two settings (Fig. 6E). GO analysis of these shared genes highlighted terms related to neurogenesis and synapse formation (Supplementary fig. 6A), which is consistent with the cellular make-up of the COs at the timepoint considered.

**Figure 6:**
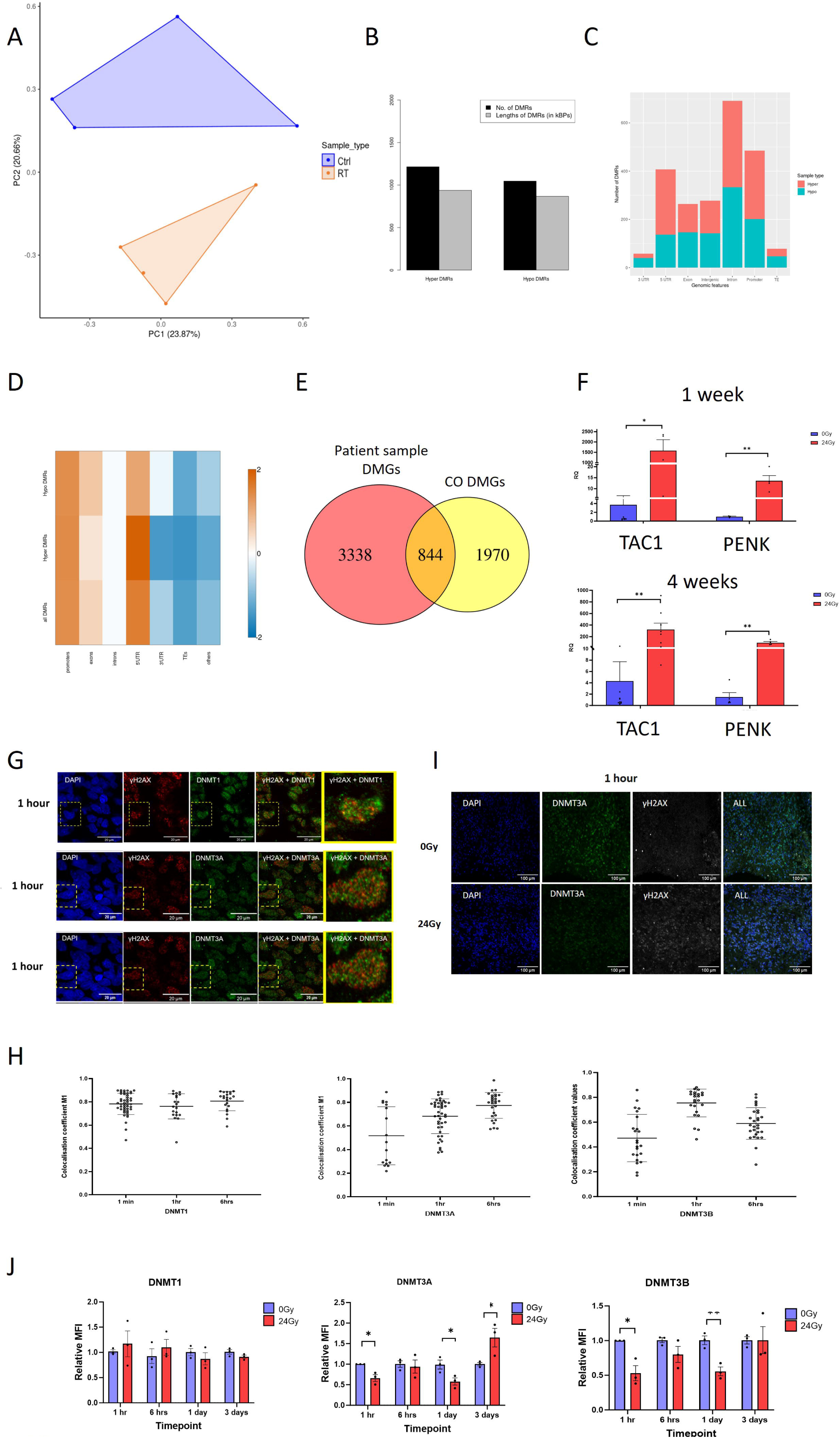
After irradiation in a CO model, there is disruption of DNA methylation patterns accompanied by upregulation of TAC1 and PENK as well as dysregulation of DNMTs. **A** PCA of DNA methylation data using all probes after standard filtering. **B** Bar plot showing number and length (in kilo-base pairs) of hyper- and hypomethylated DMRs. **C** Bar plot depicting the number of DMRs by genomic region, with proportion of hypermethylated (red) and hypomethylated (blue) DMRs. **D** Enrichment heatmap showing genomic location of DMRs, for all DMRs, hyper- and hypomethylated DMRs, as compared to whole genome as represented on DNA methylation array. Scale represents logFC(genomic features in DMRs / whole genome). **E** Venn diagram showing the overlap of DMGs from patient samples and CO model. **F** Bar plot depicting relative expression of TAC1 and PENK using RT-qPCR at 1 and 4 weeks after irradiation. Significance was tested with unpaired t-test. **G** Representative immunofluorescence (IF) images illustrating colocalisation of γH2AX (red) with DNMT1, DNMT3A and DNMT3B (green) at 1 hour post-irradiation. The overlay images demonstrate points of colocalisation in yellow. Nuclei were counterstained for DAPI (blue). Scale bars are 20um. See also supplementary figure 7B-D. **H** Dot plots of Manders colocalisation coefficient (M1) value for γH2AX colocalising with DNMT1 (left), DNMT3A (centre) and DNMT3B (right) from individual nuclei from COs at timepoints ranging from 1 min to 6 hours post-irradiation. The error bars represent SD. See also supplementary fig. 7B-D. **I** Representative IF images of DNMT3A (green) and γH2AX (white) at 1 hour post-irradiation. See also supplementary figure 8. **J** Bar plots depicting relative mean fluorescence in tensity (MFI) of DMNT1 (left), DNMT3A (middle) and DNMT3B (right) at timepoints ranging from 1 hour to 3 days post-irradiation. Each data point on the graph represents pooled data from a single CO. Unpaired t-test was performed to assess statistical significance. Error bars represent the standard error of the mean (SEM). See also supplementary figure 8.

**Figure 7:**
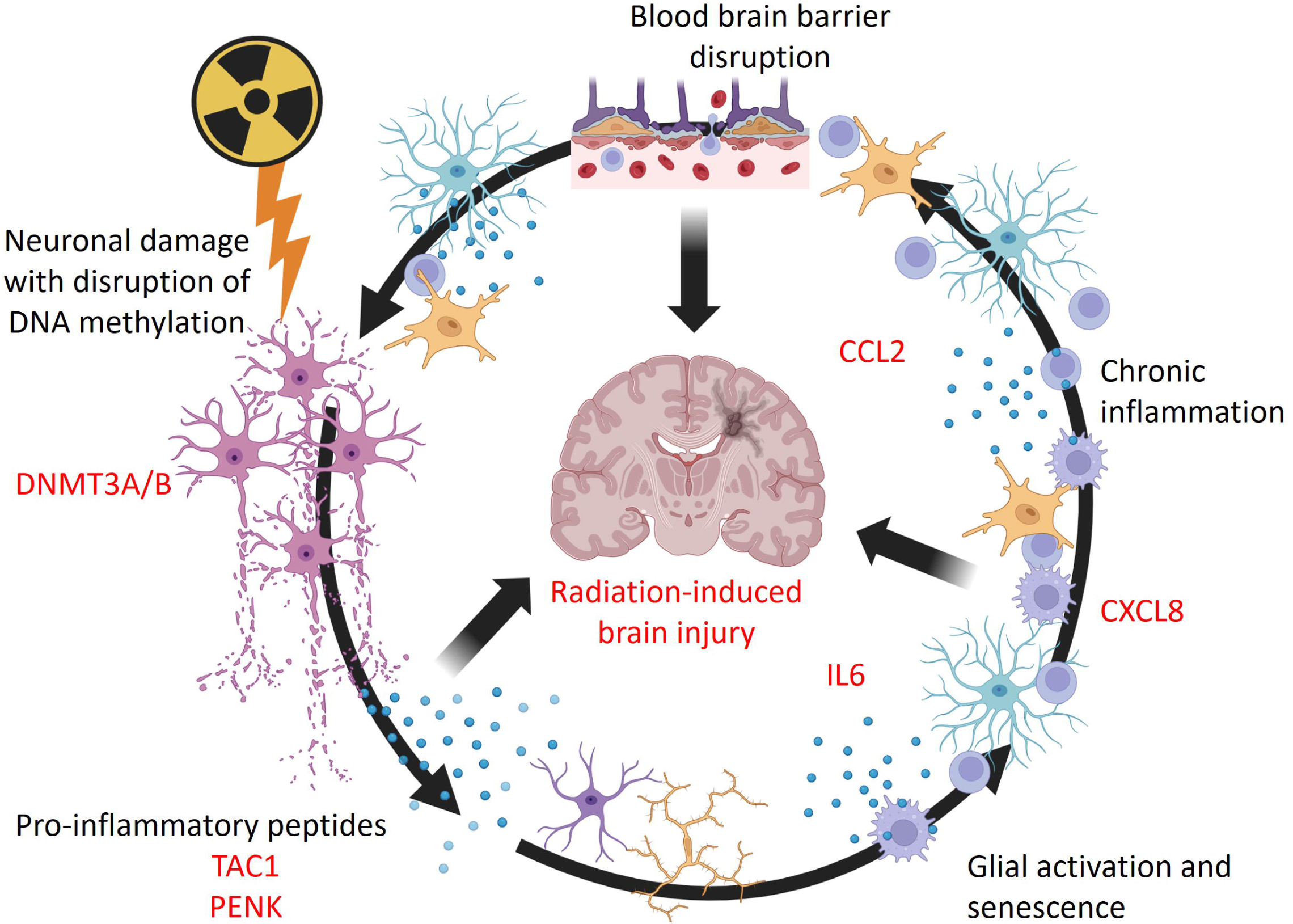
Schematic showing proposed interaction between targeted radiotherapy, disruption of DNA methylation and chronic neuroinflammation to produce radiation-induced brain injury. See main text for details; created using BioRender.

39% (330/844) of these shared DMGs had the same direction of methylation, however, given the large number of shared DMGs that we have seen here, these genes likely represent those which are highly susceptible to disruption in their methylation after irradiation in neurons. Significantly, included within these shared DMGs were both *TAC1* and *PENK* (both hypermethylated). We then tested expression levels of these two neuropeptide precursors in the CO model at two timepoints after irradiation and found that the levels of both were increased (Fig. 6F), suggesting that they are upregulated as an early response, before any downstream chronic neuroinflammation has developed.

We have shown that there is a disruption of DNA methylation in both human tissue samples and a CO model, which shares some common features, and we therefore next wanted to examine whether any of the underlying mechanisms of the DNA methylation machinery were disrupted after irradiation. DNA breaks caused by radiation occur through direct particle damage and indirect damage through the generation of ROS. Indeed, in our CO system both mechanisms were active immediately after irradiation, as assessed by DNA damage markers, up to 6 hours post irradiation, with DNA damage resolved at 1 day, consistent with established timeframes[58] (Supplementary fig. 6E). Conversely, the NF-kB pathway was not activated (there was no evidence of nuclear translocation of NF-kB p65) after irradiation in our CO model (Supplementary fig. 7A). This pathway is a common effector of cellular responses after stress and DNA damage[59], but does not appear to be significantly contributing in our model.

Whilst there have been some mechanisms proposed as to how radiation could disrupt DNA methylation, the process by which this happens at clinically relevant radiation doses is not well described. The relationship between radiation and DNA methylation changes is complex, but is thought to involve direct physical effects on the methylome mediated by ROS and dysregulation of enzymes that methylate or demethylate cytosines[7, 8], including the maintenance DNA methyltransferase (DNMT) 1 and the de novo methyltransferases DNMT3A and DNMT3B. There is evidence that these three DNMTs are involved in repair of DNA methylation patterns during the DNA damage response[60–62]. Indeed, irradiation of COs revealed colocalisation between the DNA damage marker γH2AX and DNMTs (Fig. 6G, H). DNMT1 was colocalised at all immediate timepoints where DNA damage was observed (Supplementary fig. 7B), whereas for DNMT3A this was most pronounced at 6 hrs (supplementary fig. 7C) and for DNMT3B at 1 hour post-irradiation (supplementary fig. 7D).

We then assessed the levels of DNMTs with immunofluorescence in the COs at timepoints up to three days after irradiation and found that there were decreases of DNMT3A and 3B at one hour and one day timepoints, with a subsequent increase in DNMT3A levels at three days, whilst the levels of DNMT1 were unchanged over the same period (Fig. 6I,J and supplementary fig. 8). These results suggest that dysregulation of the DNA methylation machinery is occurring in neurons at early timepoints after irradiation at clinical doses.

In a CO model system with a clinically relevant radiation dose and at early timepoints, we have shown that there is a disruption of DNA methylation patterns. This disruption has several similarities to the methylation changes observed in patient samples and there is a large overlap in DMGs between the two settings. We have also shown that two of the neuropeptides identified as potential drivers of neuroinflammation in our ST data are also upregulated in COs after irradiation. In addition, we have shown that there is dysregulation of the DNA methylation machinery occurring immediately after irradiation.

## Discussion

In this study, we have examined the DNA methylome and transcriptional landscape of a unique cohort of human brain tissue samples that had received targeted radiotherapy. Using bulk -omics techniques we have identified distinct differences in DNA methylation patterns in peri-lesional brain tissue. Further characterisation of this tissue with spatial transcriptomics identified specific micro-environmental niches present after irradiation and, within an irradiated neuronal niche, highlighted neuropeptide precursors that could be propagating neuroinflammation. We then modelled early radiation changes, focussing on those occurring in neurons, in a cerebral organoid system and compared this to human tissue samples. We show similarities in DNA methylation disruption and identified genes that are susceptible to irradiation-induced DNA methylation changes. Among these genes, selected neuropeptide precursors showed increased expression early after irradiation, supporting their role in initiating radiation-induced neurotoxicity. Finally, we demonstrate that specific elements of the DNA methylation machinery, including the DNMTs, are dysregulated after irradiation, likely linked to DNA damage repair.

The alteration of DNA methylation has been shown to be involved in the pathogenesis or disease progression of many neurological disorders, including neurodegeneration[63], multiple sclerosis[64], and stroke[65]. There are many studies that have shown that radiation can also disrupt DNA methylation patterns, however, the findings are highly variable, dependent on type of radiation, radiation dose, and on the animal model or cell type used. Additionally, the evidence for these epigenetic alterations is derived almost entirely from neoplastic cells and non-CNS mouse tissues, with no investigations in the human brain tissue collected in a clinically relevant setting. The distinct DNA methylation changes we have shown included a bias towards hypomethylation of DMRs, with DMRs preferentially found in promoter, exon and 5’UTR regions, suggesting a strong influence on expression as well as context-specific target-gene hypermethylation. Whilst these findings are in keeping with the current coarse description in other tissues/cells [9, 66], this is the first time a detailed characterisation of the DNA methylation changes in patient tissue after radiotherapy has been performed.

The cohort in this study had a time between radiotherapy and subsequent tissue resection of 7-240 months, ensuring that we were able to study changes in a timeframe that is meaningful for patients in which the long-term sequelae of radiotherapy occur, however, we are unable to capture the immediate changes after radiation in these samples. Radiotherapy is a potent inducer of DNA damage, and it has been shown that stable changes in DNA methylation after DNA damage can occur within a discrete early timeframe[62] and that this early disruption could persist as epigenetic ‘scars’[62, 67, 68]. Supporting this, several animal models have shown disruption of DNA methylation that persists in body organs for months after radiation exposure[69–72], whilst many of the side-effects of radiation are observed up to years after treatment, suggesting a possible link between DNA methylation alterations and radiation-induced late side-effects. Since the common consensus is that neuronal damage is a major driver of radiotherapy induced neuroinflammation, we used a cerebral organoid (CO) model to capture early changes that could be leading to these epigenetic scars in neurons. Indeed, when we examined the DNA methylome at early timepoints after irradiation in COs we again found distinct patterns of DNA methylation and that DMRs were distributed similarly across the genome as in patient samples. Compared to the DMRs identified from patient samples, those seen in the CO were more hypermethylated overall. This difference may be because within our patient tissue samples we are capturing a picture of the DNA methylome from a more diverse cell population, and it is likely that the chronic inflammatory environment in which these cells are situated has an effect on DNA methylation status. Therefore, we are arguably capturing neuronal and irradiation specific DNA methylation changes in our CO model.

Interestingly, when we compared genes that were differentially methylated in patient tissue samples and in COs after irradiation, we identified a large number of genes common to both settings. These genes may represent those that are particularly sensitive to DNA methylation disruption, both immediately after irradiation and which persists over time. The response of neurons to DNA damage is complex, with repair of DNA damage not uniform across the genome, but dependent on cell-specific transcriptional activation and other factors[73, 74]. There are multiple lines of evidence linking the reconstitution of DNA methylation patterns and DNA damage repair pathways[61, 75]. Whilst the exact mechanisms of DNA methylation reconstitution in this context are not completely understood in human tissue, it appears that this process is error prone. These errors can lead to a revision of the methylation landscape in repaired genes, producing ‘epi-alleles’ in individual cells that lead to differential gene expression[62, 67, 76].

Additionally, in this study we have shown there are changes in the levels of DNMT3A and DNMT3B at early timepoints after irradiation. These DNA methyltransferases have been shown to be involved in DNA methylation reconstitution[60–62, 77], and we have seen their colocalisation with DNA damage foci in our CO model. This colocalisation is occurring in the same timeframe at which the changes in DNMT levels are observed, therefore, their dysregulation at these early timepoints will also lead to disruption of DNA methylation after the DNA damage caused by irradiation. We see a decrease of DNMT3A and DNMT3B at one hour and one day, after which DNMT3B levels return to baseline but DNMT3A levels subsequently increase at three days, suggesting that there are complex dynamics at play and that further work is warranted to explore how this may affect DNA methylation at global and gene specific levels.

All the above factors are likely combining after irradiation to manifest as a susceptibility of specific genes to disruption of DNA methylation after the DNA damage inflicted by radiation. Thus, if the disruption of DNA methylation causes prolonged upregulation of a pro-inflammatory gene, then this could initiate a persistent increased inflammatory response. Unresolved (chronic) inflammation is characterised by the secretion of cytokines that maintain inflammation and produce redox stress, which also induces DNA damage[52]. This would further disrupt the DNA methylome in specific locations, that would further increase the initial vulnerability and so on. This feedforward process could partly explain the chronic inflammatory changes that we find in the brain after radiotherapy.

Using ST we have identified specific micro-environmental niches within the irradiated brain, and by examining an irradiated neuronal niche we identified neuropeptide precursors that: are differentially methylated in both patient tissue and COs after irradiation; are also overexpressed in irradiated neurons in both settings; and appear to be involved in active signalling in patient tissue. The *TAC1* gene encodes the tachykinins substance P (SP) and neurokinin A (NKA). Tachykinins are a family of evolutionarily preserved small neuropeptides[78] that act as neurotransmitters, but also at the intersection of immune and nervous system cellular communication[78, 79]. Substance P is known to be involved in the brain’s inflammatory response to many insults, where it induces vasodilation, promotes the recruitment of immune cells, and augments the inflammatory responses of both infiltrating and resident cells[80]. *PENK* and *PDYN* are precursors for the opioid peptides enkephalin and dynorphin, respectively. These are best known to act as neurotransmitters involved in pain processing. Additionally, opioids and their receptors are present on both glia and peripheral immune cells[81, 82]and cross-talk can occur between the opioid receptors and the chemokine and chemokine receptor families[83]. Our results suggest that the chronic overexpression of these neuropeptides could be one of the factors driving inflammation. There are drugs available that target tachykinin pathways, which have been developed to reduce neuroinflammation[80], and their use as an adjuvant after radiotherapy could now also be considered. Intriguingly, the tachykinin receptor NK1R (neurokinin type-1 receptor) has been shown to be expressed in some gliomas[84, 85], and there is evidence this pathway is also involved in brain metastasis in breast cancer[86], suggesting that treatments modulating tachykinin function could be harnessed for a dual purpose. Pharmacological targeting of opioid pathways using commonly available drugs has also been suggested as way to modulate neuroinflammation[87] and could also be utilised in the setting of radioprotection.

In conclusion, our study links radiotherapy induced neuroinflammation with disruption of DNA methylation for the first time and suggests possible driving mechanisms for this neuroinflammation. It also demonstrates that examining neurological diseases in patient tissue samples complemented with patient-derived 3D models is critical for clinically relevant advances in our understanding of complex pathological mechanisms.

## Methods

### Study cohort

Tissue samples were identified by medical records searches and slide review by a neuropathologist. We retrospectively identified formalin fixed paraffin embedded (FFPE) neurosurgical samples from patients that had undergone supratentorial targeted radiotherapy, followed by resection and that contained peri-lesional brain tissue within 15 mm of the irradiated lesion, that would have been irradiated as part of the treatment field in our centres, as confirmed by clinical oncologist. Characteristics of all patients included in the final analysis, whose samples yielded suitable quality and quantity of DNA and RNA, are summarised in Supplementary figure 1A. Ethical approval was secured for the use of all patient samples (BRAIN UK Ref: 20/005).

### DNA and RNA extraction from FFPE tissue

H&E and subsequent unstained sections were attained from the representative FFPE blocks. Concentrations and quality of DNA and RNA were quantified using the Nanodrop 1000. DNA methylation array was performed by UCL Genomics, UK and RNA transcriptome analysis by Oxford Genomics, UK. For DNA methylation, oxidative bisulfite chemistry and bisulfite conversion was applied alongside Infinium Methylation EPIC (850K) BeadChip Array for the detection of 5-methylcytosine (5mC). Transcriptome analysis involved conversion of selected mRNA to cDNA. The prepared libraries are size selected and multiplexed before 150bp paired end sequencing on the Illumina platform (NovaSeq6000). A minimum concentration of 500 ng for genomic DNA in 45 μl and 200 ng for RNA in 12 μl with 260/280 and 260/230 ratios over 1.80 was used. Samples that failed to meet these criteria were selected for further concentration and purification.

### DNA methylation analysis

Illumina Infinium Methylation EPIC array was used to investigate differentially methylated genes between experimental and control cases to assess whether DNA methylation changes occurred as a consequence of irradiation. DNA methylation profiles were obtained from FFPE human brain tissue and COs. Firstly, the data quality was assessed, and standard filtering of the EPIC probes was performed to filter out any failed probes, low-quality probes, and cross-reactive probes. EPIC array CpGs in the sex chromosomes were disregarded to reduce the bias. For the normalisation step, the wateRmelon package was used to normalise the beta values of the remaining probes. In addition, probes relating to single-nucleotide polymorphism (SNPs) were disregarded. Differentially methylated probes (DMPs) and differentially methylated regions (DMRs) were identified using the R package DMRcate, with the FDR value set to 0.01 for identification. Differential analysis was performed to obtain DMPs and DMRs by comparing the irradiated samples with the control. To qualify as a DMR, the criteria were that the DMRs should contain at least 6 CpG sites and be separated by no more than 400 base pairs. All analyses and manual DMR annotations of different genomic features and their corresponding enrichment analysis in DMRs were carried out based on the Ensembl annotation of hg38, v110. Hierarchical clustering was performed using the R package ComplexHeatmap and 2D/3D principal component analysis with the R package ggbiplot to perform exploratory analysis. The analysis was conducted to understand preliminary relationships between samples and reveal underlying data patterns based on different treatment conditions, i.e. irradiation. Further analysis was performed to determine the genome regions affected. Differentially methylated genes were therefore acquired for subsequent analysis.

### RNA sequencing analysis

Regarding RNA sequencing analysis, the transcriptome profile of FFPE human brain samples was analysed. The raw RNA-seq data was examined using FastQC v0.12.14 and thereafter trimmed using TrimGalore v0.6.105 . Alignment to the reference genome (Homo_sapiens.GRCh38.cdna.all.fa.gz, release 110) was performed using Salmon v1.10.2 6. R package trimeta 7 was utilised to read and normalise the raw counts for different samples. This was followed by conversion of the normalised data from the transcript level to the gene level. Genes that have no counts or only one count across the samples were filtered out. The converted normalised raw counts were further converted to counts per million (CPM) and genes reporting 0 CPM were filtered. The differential expression analysis (control versus experimental) was carried out using the package DESeq2. Exploratory analysis was carried out, including hierarchical clustering and differential analysis was performed to acquire differentially expressed genes by comparing the irradiated samples with the control. Ultimately, a list of differentially expressed genes was acquired for further analysis. Genes with significant differential expression were selected based on the adjusted p-value (p.adj < 0.05).

### Integration and comparative analysis

Integration of methylome and transcriptome data was performed to identify concordant genes to understand whether methylation impacts gene expression and to delineate epigenetically mediated changes in the expression profile. Genes that demonstrated hypermethylation/downregulation and hypomethylation/upregulation were considered. Comparative analysis was also performed between the FFPE brain vs COs for the DNA methylation data to ascertain the shared and consistent methylation marks.

### Pathway analysis

The most significantly differentially expressed or methylated genes (see figure legends for details) were submitted for Gene Ontology (GO) analysis using g:Profiler[88] with g:SCS significance threshold and pathway p-value threshold of 0.05. Pathway analysis was run for Gene Ontology of Biological Processes and Molecular Functions, with no electronic annotation, and .gem and .gmt files were exported and visualised on Cytoscape_v3.10.1 using the EnrichmentMap app. Pathways were manually grouped and annotated. Nodes are coloured by FDR value and size is proportional to the number of genes included in each term. Edge thickness represents similarity between genesets. Edge cutoff was set at 0.5 unless otherwise stated. Gene level association was performed using String[89] and visualised in Cytoscape_v3.10.1 using the String app, with clustering performed with a granularity value of 4.

### Spatial transcriptomics

Case numbers used and numbers of spots assessed were comparable to published literature[30, 39, 90, 91]. Spatial transcriptomics (ST) was conducted using 10X™ Visium Spatial Gene Expression Slide & Reagent Kit, 16 rxns (PN-1000184), according to the protocol detailed in document CG000239 available in 10x demonstrated protocols. 10 micron-thick tissue sections were mounted on the ST slides and stained with H&E described in document CG000160 available in 10X demonstrated protocols. Imaging of whole slides was done at 20X magnification on a Nikon Eclipse 80i Stereology Microscope. The remaining steps were conducted according to the manufacturer’s protocol. The libraries were sequenced on multiple Illumina Nextseq 2000 (paired end dual-indexed sequencing) flowcells to achieve the recommended number reads per ST spot. The ST samples were prepared using 10X genomics Space Ranger software, which uses tiff image files of tissue, raw fastq sequencing files and 10X Visium slide information to align raw reads to the reference genome and output count matrices and annotated tissue section images. The reference genome used for alignment was builts from Ensemble GRCh38 version 91 assembly. All other parameters for generating the counts data for ST were set to default settings. Slides with overlying spots were visualised in Partek Flow (Illumina). CPM normalisation was performed and after removal of any spots containing tumour cells, by neuropathological assessment and computational filtering for tumour cells that had not been removed manually. PCA was performed for dimensionality reduction, and output used for unsupervised clustering using the SLM method with default settings other than a resolution of 0.75 and PC of 5. Unsupervised clustering was then visualised with 2D UMAP with cluster annotation in Partek Flow. Bubble plots were generated in Partek Flow using the hierarchical clustering / heatmap function, using normalised count matrices. Differential expression analysis between clusters was performed using the GSA algorithm in Partek Flow using default settings. Spatial plots with overlying clustered spots in supplementary figure 2A were generated in Partek Flow using the Visium plot function.

### Signature scores

Signatures were extracted from the cited literature uploaded to Partek Flow using the List function. Scores for each spot were calculated in Partek Flow. Scores were then normalised for gene count per spot and truncated violin plots produced using GraphPad Prism and significance calculated with unpaired t-test.

### Reference-free deconvolution

Spatial deconvolution of tissue spots was performed using the STdeconvolve package[26]. In brief, h5 files and spatial images and coordinates produced by the 10X Space Ranger pipeline were used as inputs. Samples spots were filtered as above and count matrices and tissue spot positions were then extracted from each sample. Further filtering was performed to remove poor barcodes and features using the stDeconvolce cleanCount function, the following parameters were used: min.lib.size = 250, min.reads = 10, min.detected = 5. Next the top 500 over-dispersed genes in each dataset were identified and then the common over-dispersed genes shared by all samples were identified. All sample count matrices were then merged into a single count matrix based on common over-dispersed genes. LDA models were k varying between 2 and 18 were fitted to the merged count matrix. The optimal k was found to be 9 and results were plotted using stDeconvolve functions. Markers of each potential cell type were identified to determine the identities and finally transcriptional profiles of each of the potential cell types compared. Cell types were annotated using a combination of the most highly expressed genes, using the Human Protein Atlas and STAB database[29], and the spatial location, confirmed by a neuropathologist.

### Correlation heatmap

To group similar clusters, we computed cluster gene specificity scores for all cluster marker genes (mean normalised counts per cluster/total mean normalized counts) – and performed Pearson correlation between cluster gene specificity signatures, representing data as a heatmap.

### Proportional differences

Proportional differences in fig. 3G were calculated using an Rlibrary to calculate the proportional difference in cell number proportions between clusters in scRNA-seq, available at https://github.com/rpolicastro/scProportionTest/releases/tag/v1.0.0[92].

### Receptor-ligand interactions

Two approaches were adopted to perform spatial recpeot-ligand interaction analysis. First, custom code was written in order to compare the interaction scores of receptor-ligand pairs between clusters, based on the work from Kaufmann et al.[39]. In brief, the expression of each receptor and ligand, listed in the CellPhoneDB[44] database, in each tissue spot was determined. In the case of receptor complexes the expression value of the least expressed subunit was taken as the expression value. If any subunit was not expressed the expression value of the complex was set as 0. Next interaction scores were determined for each tissue spot, and its immediate surrounding neighbours. Interaction scores were also calculated within a tissue spot. The interaction score was defined as the mean expression of receptor A and ligand B in tissue spot 1 and tissue spot 2 respectively. Receptor expression was always taken from spot 1 – therefore spot 1 was the receiver of signalling, whilst ligand expression was always taken from spot 2; therefore spot 2 was the sender of signalling. This calculation processes was iterated across all tissue spots in combination with all neighbours of each tissue spot. Once interaction scores were calculated each tissue spot was annotated with its cluster identity. In this way, the interaction scores for a given cluster could be extracted. Extracting interaction scores based on the spot 1 cluster annotation identified the interaction scores that spots for a given cluster were *receiving*, whilst using the spot 2 cluster annotation identified the interaction scores that spots for a given cluster were *sending*, because receptor expression was always taken from spot 1 whilst ligand expression was always taken from spot 2. Finally, interaction scores for each receptor-ligand pair detected were compared between two clusters, in terms of both what each cluster was sending and receiving. Interaction scores between clusters were compared using a Wilcoxon Test and p-values adjusted for multiple hypothesis testing using the Benjamini-Hochberg procedure. All code produced for this spatial neighbourhood analysis has been deposited on Github (https://github.com/jamesboot/interactVis). Second, the CellChat v2 package[49] was also used to perform spatial receptor-ligand interaction analysis. In brief, 5 samples (RT14, RT13, RT6, RT1 and Cnt8) were analysed for significant receptor-ligand interactions between their cluster annotations. As previously, h5 files and spatial images and coordinates produced by the 10X Space Ranger pipeline were used as inputs. Raw data were filtered and normalised, and then samples analysed, and results visualised as per CellChat v2 instructions. Finally results for pathways of interest were assessed and visualised at both the sender and receiver level.

### Cerebral organoid culture

Cerebral organoids (COs) were established from EPSCs derived from fibroblasts of dura mater from two different patient lines: DURA19 and DURA61. When the cells reached 70% confluence, they were harvested for the development of COs. The number of viable cells was calculated using a haemocytometer and trypan blue staining. 70,000-80,000 EPSCs were used for the development of individual organoids using the STEMdiff™ Cerebral Organoid Kit (Stemcell technologies, #08570) and the STEMdiff™ Cerebral Organoid Maturation Kit (Stemcell technologies, #08571), following the producer’s protocol. Briefly, the cells were transferred into embryoid body (EB) media to aid the development of embryoid bodies in round bottom ultra-low attachment 96 well plates (Costar, #7007). During the fifth day of the development process, when the EB diameter had grown to a minimum of 300µm, the EB bodies were cultured in 24-well ultralow attachment plates (Corning®, #3473). STEMdiff™ Cerebral Organoid Basal Medium 1 and Supplement B were used to initiate the development of the neural ectoderm layer. ROCKi was also incorporated in the media during the first four days of culture. Once neuro ectoderm development was successful, the EBs were embedded in Matrigel droplets on day 7 and cultured in neural expansion media to encourage the neuroepithelial bud expansion. Parafilm (Sigma-Aldrich, # HS234526B) was sterilised and used as embedding surface. Circular indentations were created on a sterilised parafilm contained in a 10mm dish. Each EB was placed onto individual indentations and any surplus media was removed before adding Matrigel (Corning®, #354277). The EBs are immersed in Matrigel and incubated at 37⁰C to allow the gel to set. Next, the structures were carefully washed off from the parafilm with neural expansion media (STEMdiff™ Cerebral Organoid Basal Medium 2, Supplement C and Supplement D) and transferred onto 6-well ultralow attachment plates (Corning®, #3471), with approximately ten EBs per well. Finally, the organoids were cultured in CO maturation media (STEMdiff™ Cerebral Organoid Basal Medium 2 and Supplement E) on day 10 and transferred to an orbital shaker at 100 RPM for the remaining culture period to further aid neural tissue growth and development. The organoids underwent media change every 3 to 4 days in the maturation media and reached maturity on day 30. Irradiation was performed using X-Ray Biological Irradiator within the Queen Mary Biological Science Unit and radiation dose of 24 Gy was administered at 8.5Gy/min.

### DNA and RNA extraction from COs

COs were dissociated. The RNA/DNA/Protein Purification Plus Kit (Norgen, #47700) was utilised to extract DNA and RNA, as per the manufacturer’s instructions. For the extraction of DNA and RNA when working with organoids embedded in gelatine, extra wash steps were undertaken before processing.

### Histological and immunofluorescence staining

Hematoxylin and eosin (H&E) staining was performed using standard protocols and slides were imaged using the EVOS XL core imaging system for brightfield images. Immunofluorescence staining was performed using standard protocols. The following primary antibodies were used in the CO characterisation study: goat anti-Sox2 (1:500, Santa Cruz), rabbit anti-NeuN (1:200, Abcam), mouse anti-Nestin (1:200, Chemicon), rabbit anti-TBR1 (1:200, Abcam) and mouse anti-TUJ1 (1:200, Abcam). For irradiation experiments, the following antibodies were utilised: mouse anti-phospho-Histone-H2AX (1:250, Merck), mouse anti-8-Hydroxy-2’-deoxyguanosine (1:100, Abcam), rabbit anti-NF-kB p65 (1:100, Abcam), rabbit anti-DNMT1 (1:100, Abcam), rabbit anti-DNMT3A (1:200, Cell Signaling) and rabbit anti-DNMT3B (1:200,Abcam). The following secondary antibodies were used in the study: donkey anti-goat antibody Alexa Fluor 647 (1:200, Invitrogen™), donkey anti-rabbit Alexa Fluor 546 (1:200, Invitrogen™), donkey anti-mouse Alexa Fluor 488 (1:200, Invitrogen™), goat anti-rabbit Alexa Fluor 546 (1:200, Invitrogen™), goat anti-mouse Alexa Fluor 488 (1:200, Invitrogen™), goat anti-mouse Alexa Fluor 546 (1:200, Invitrogen™), goat anti-rabbit Alexa Fluor 546 (1:200, Invitrogen™), goat anti-mouse Alexa Fluor 647 (1:200, Invitrogen™) and goat anti-rabbit 647 (1:200, Invitrogen™).

### Confocal microscopy and quantification

Immunofluorescence images were acquired with the Zeiss 710 Laser Scanning confocal microscope. Images were captured with 20X and 63X magnification lenses depending on the nature of staining, and a series of Z-stack images were acquired for those that required assessment of signal dynamics. For images that required quantification, confocal calibration settings remained consistent across each staining set. Images were then quantified and processed in FIJI ImageJ. The region of interest was designated by creating a mask on the nuclei staining to extract the average intensity value. The mask threshold was kept automatic to distinguish between the background and the signal. The mean grey value was then calculated from the applied mask corresponding to the signal of interest. Five to ten images per sample, form three samples per condition, were captured for quantification. The relative mean fluorescence intensity (MFI) value was subsequently calculated using Microsoft Excel. To quantify colocalisation, the Manders’ colocalisation coefficient (M1) value was calculated to determine the fraction of overlap between two signals. Manders’ coefficient is a well-established measure which calculates the co-occurrence fraction of a fluorescent signal from one channel with that of another. M1 value ranges from 0 to 1, with 0 indicating total absence of colocalisation and 1 indicating perfect colocalisation[93]. Just Another Colocalization Plugin (JACoP) from Image J was used to carry out the M1 calculation from the region of interest. A series of Z-stacks containing 8-12 images were processed at 63X magnification to calculate the colocalisation between signals in the set of images. The background was subtracted, and the noise was reduced with the despeckle tool to ensure the exclusion of artefacts from the calculation. Following the selection of nuclei of interest, z-stack images corresponding to these nuclei were determined, and the region of interest was isolated using the crop tool. The JACoP plugin created an automatic mask from the two selected channels of interest, and the M1 value was calculated, generating an overlap value from the signals analysed.

### RT-qPCR

SuperScript III Reverse Transcriptase (Invitrogen™, #18080093) was used for the reverse transcription process as per the manufacturer’s protocol. Random primers were mixed with RNA and diluted with PCR grade water, making up to 10 µL of solution. Primers were annealed at 65°C for at least 1 minute and then cooled at 4°C for 5 minutes. 5X FS Buffer, 0.1 M DTT and SuperScript III Reverse mix was added to the solution. The mixture was then incubated at 25°C for 5 minutes and incubated at 50°C for 30 minutes. Reaction time is increased to 55°C thereafter and finally, the inactivation step is carried out by heating at 70°C for 15 minutes. The cDNA was diluted to a concentration of 2.5 ng/μl using PCR grade water and stored in -201C until further qPCR reactions were carried out. For the qPCR reaction, predesigned primers KiCqStart® SYBR® Green Primers (Sigma-Aldrich, # KSPQ12012) were used for gene expression analysis. To prepare the qPCR mix, add 5 μl of the forward and reverse primers with per 90 μl of PCR grade water for a final concentration of 10 μM. 96-well PCR plates (Applied Biosystems™, #4346906) were used to carry out the qPCR reaction with each well comprising of 1μl of the forward and reverse primer mix, 2 μl of diluted cDNA (5ng), 6 μl of PowerUp™ SYBR™ Green Master Mix (Applied Biosystems, # A25742) and 3 μl of PCR grade water. TAC1: forward primer (ATTCTGTGGCTTATGAAAGG), reverse primer (CATTGACACAAATGAAGCTG). PENK: forward primer (AACTGTCATTTCAGGTTCTG), reverse primer (TTTATGCACTTGGGGTAATAG). StepOnePlus™ Real-Time PCR System and StepOne Software (v2.3) were used to run the qPCR reaction. In addition, fast Cycling Mode (Primer Tm ≥60°C) was selected from the PowerUp™ SYBR® Green Master Mix manual for the qPCR thermal-cycling conditions. Cycle threshold (Ct) values were obtained from each gene run within a single reaction. ACTIN was designated as the housekeeping gene, and the cycle threshold (Ct) values for the gene were recorded with up to 40 cycles run. The Ct from genes of interest were normalised to that of the housekeeping gene. Subsequent calculations were performed in Microsoft Excel to attain the ΔCt values from Ct(gene of interest)−Ct(housekeeping gene). ΔΔCt values were derived by calculating the ΔCt difference between the irradiated samples and the control. The Relative quantity (RQ) of gene expression was calculated using the 2^(−ΔΔCt). GraphPad Prism 9 was used to visualise data.

### Statistical analysis using GraphPad Prism v9

Normalised signature scores, colocalisation, RT-qPCR, and immunofluorescence values were imported into GraphPad Prism version 9 for statistical analysis and visualisation. Unpaired t-tests were used to compare between two datasets, and analysis of variance (ANOVA) was carried out to investigate more than two groups. P-values for statistically significant results are represented in the graphs are as follows: *= p ≤ 0.05, **= 0.001 < p ≤ 0.01 = **, and ***= p ≤ 0.001, ****= p ≤ 0.0001. Error bars presented standard error of mean (SEM) or standard deviation (SD) depending on the nature of the analysis.

## Author contributions

T.O.M., P.P. and X.Z. designed and performed experiments and analysed results. T.O.M., Y.X., J.R.B. and J.N. designed and performed computational analyses. Z.A. and N.R. performed FFPE tissue sectioning. P.S. performed Visium slide processing. C.M. provided essential expertise. E.M., G.M. and D.P. provided tissue samples and essential expertise. A.W.M. and N.K. provided tissue samples. R.L. identified patients and provided essential expertise. S.B. provided essential expertise and supervision. S.M. and T.O.M. conceived the study and secured financial support for the project. T.O.M. wrote the manuscript with contribution from all authors.

## Acknowledgements

This work is funded by grants from NIHR (CL-2019-19-001 to T.O.M.), The Pathological Society of Great Britain & Ireland (JCLSG 1021 01 CL support grant to T.O.M.), Barts Charity (MGU0447 programme grant to S.M.), Brain Tumour Research (Centre of Excellence award to S.M.) and Cancer Research UK (C23985/A29199 programme award to S.M.). Ethical approval for use of human tissue samples was attained through BRAIN UK (Ref: 20/005).

---

**Supplementary figure 1:**
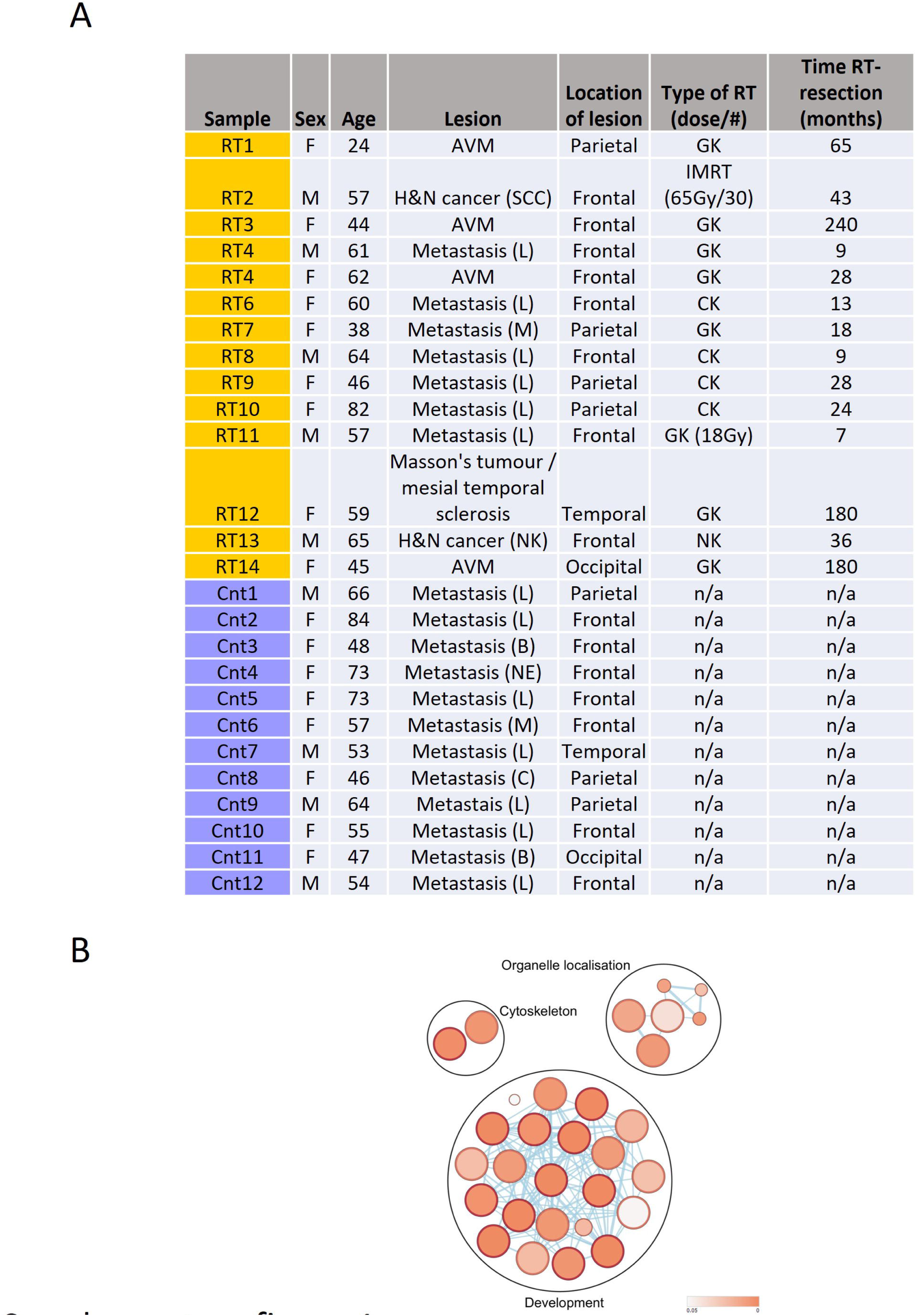
**A** Patient sample clinical characteristics. AVM - arteriovenous malformation; B - breast primary tumour; C - cervical primary tumour; CK – CyberKnife; GK – GammaKnife; IMRT - intensity-modulated radiation therapy; L - lung primary tumour; M – melanoma; n/a – not applicable; NE - neuroendocrine primary tumour; NK - not known; RT – radiotherapy; SCC - squamous cell carcinoma primary tumour. Dose is routine targeted radiotherapy dose (∼24Gy) in single fraction unless specifically stated. **B** GO analysis using all (159) concordant genes from DMG and DEG overlap.

**Supplementary figure 2:**
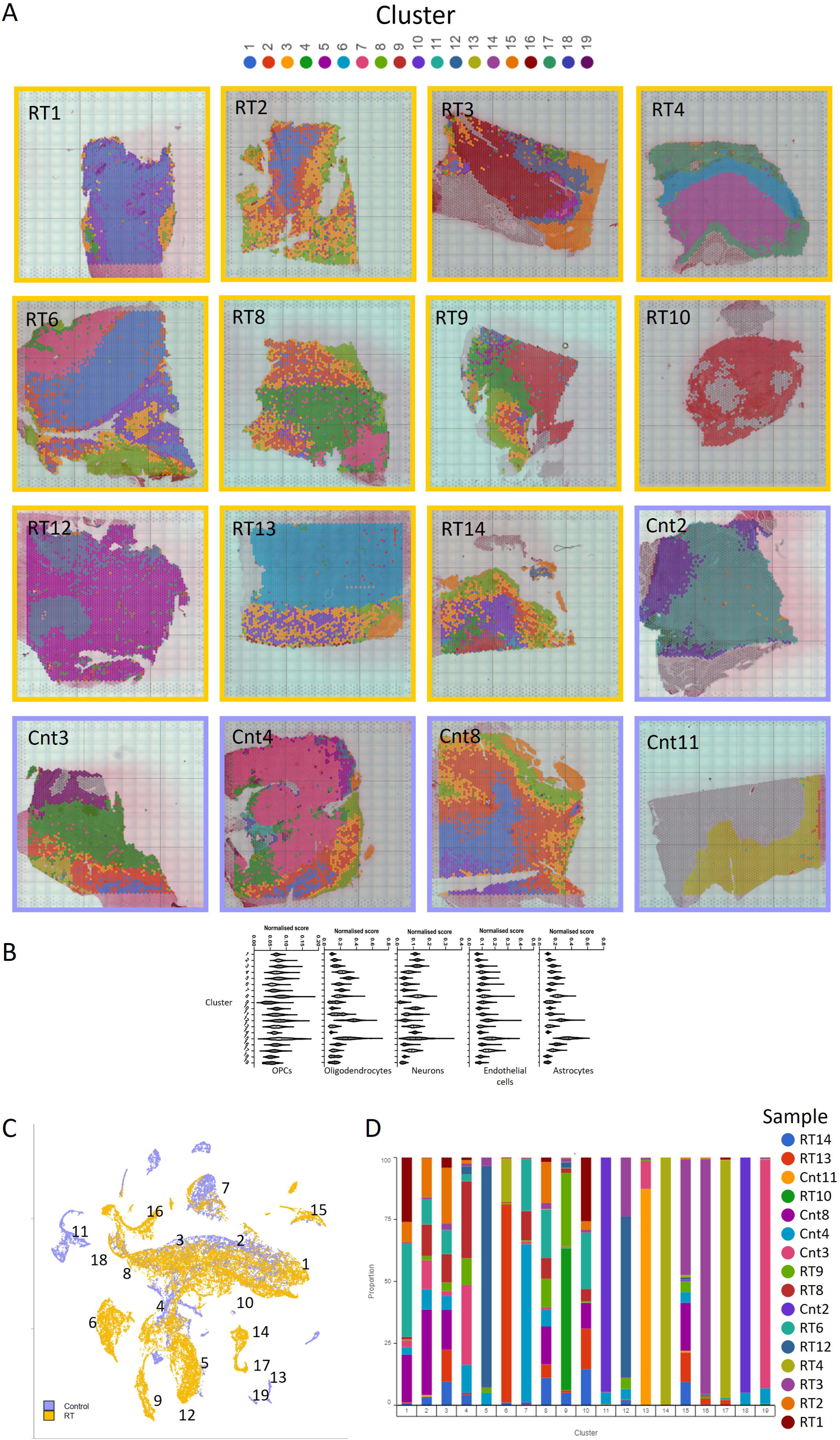
**A** Spatial transcriptomic slide overviews with spots overlying histology images. Spots are coloured by cluster, as in Fig. 3A. Border colour represents experimental group as in Fig. 1A. Dimensions of fiducial frame are 6.5 x 6.5 mm. Spots are no to scale. **B** Violin plots for cell type signature scores (from [30]) for each cluster. Y axes show z-score normalised against gene expression per spot (violin colours correspond to the colours of the clusters in Fig. 3A). **C** UMAP plot showing all ST spots. Each dot corresponds to a single ST spot, coloured by irradiated or control. Unsupervised cluster numbers are shown, as in fig. 3A. **D** Bar plot showing proportion of spots by sample for each cluster.

**Supplementary figure 3:**
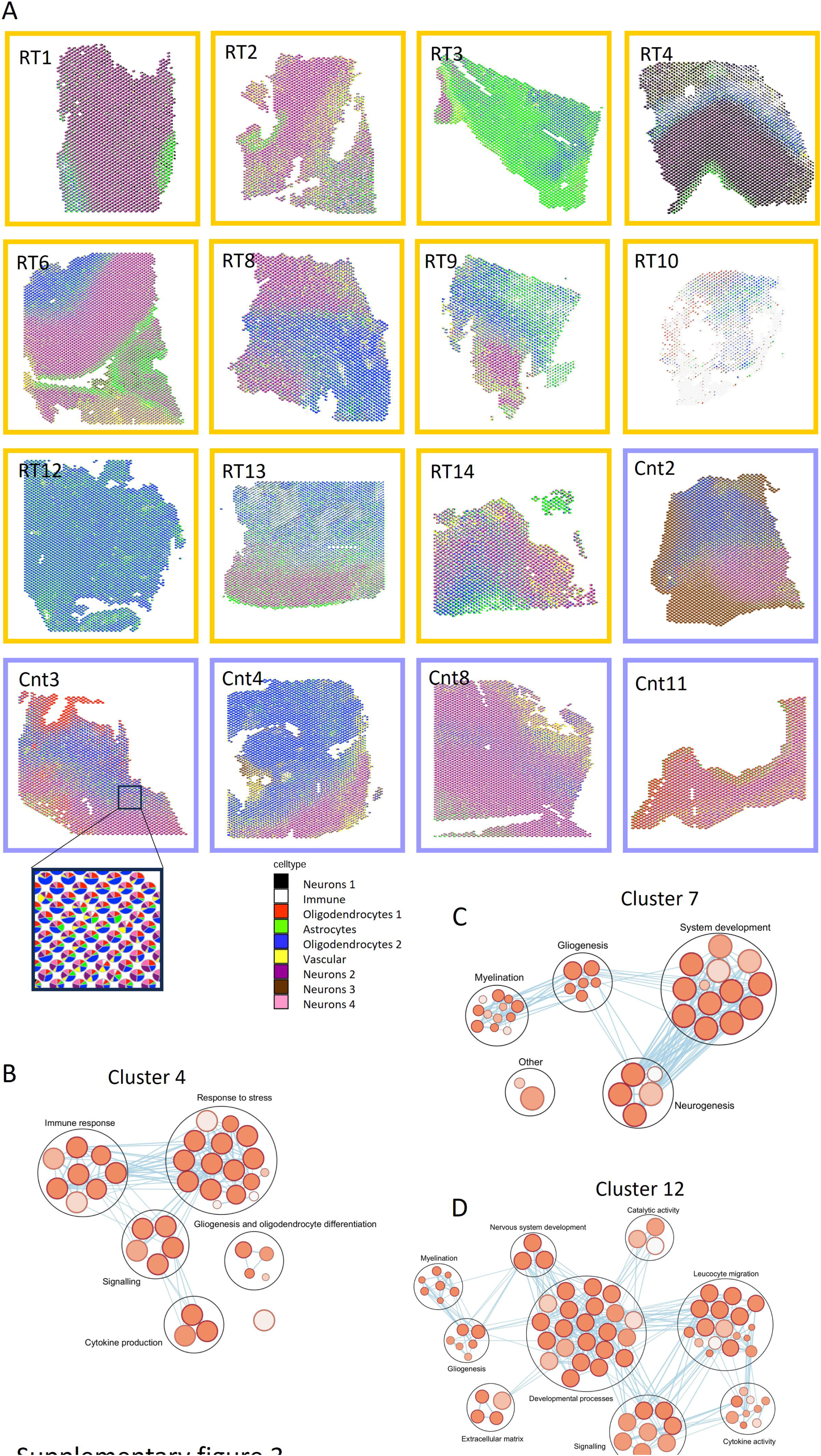
**A** Deconvolution of ST spots using STdeconvolve. Spot composition is visualised as a pie chart (shown enlarged in inset) and projected on the spatial coordinates. **B, C, D** GO analysis using top 150 cluster marker genes for cluster 4 (**B**), cluster 7 (**C**) and cluster 12 (**D**).

**Supplementary figure 4:**
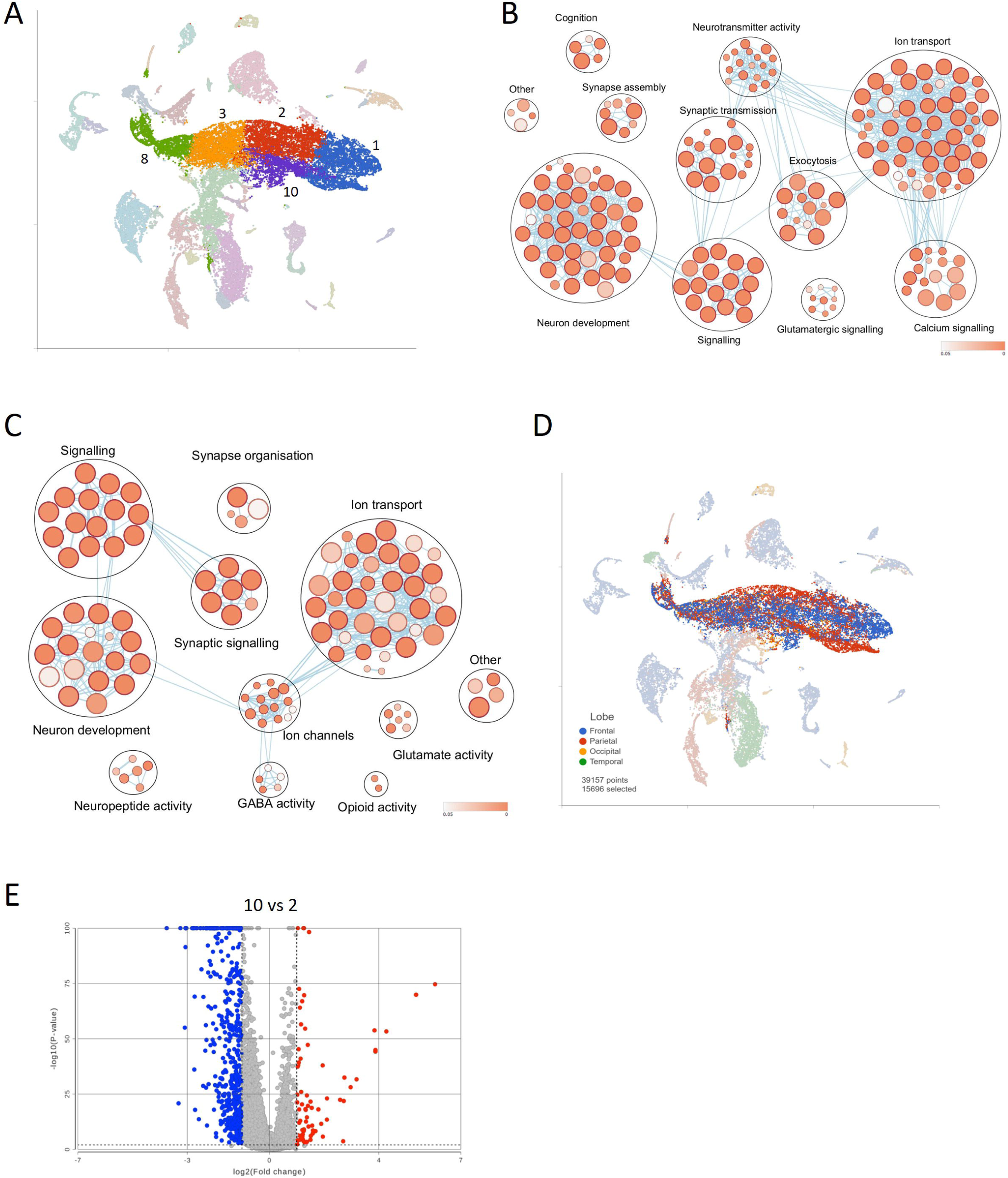
**A** UMAP as in figure 3A. Dots coloured by cluster with neuronal niches highlighted. **B** GO analysis using top 150 cluster marker genes for cluster 2. Edge cutoff 0.6. **C** GO analysis using top 150 cluster marker genes for cluster 10. Edge cutoff 0.6. **D** UMAP as in figure 3A. Dots coloured by brain region of sample. **E** Volcano plot showing differentially expressed genes between cluster 10 (irradiated glial) and cluster 2 (non-irradiated glial). Blue dots are significantly overexpressed in cluster 12 and red are significantly overexpressed in cluster 7. Significance threshold is -2.0<FC>2.0 and p-value <0.01.

**Supplementary figure 5:**
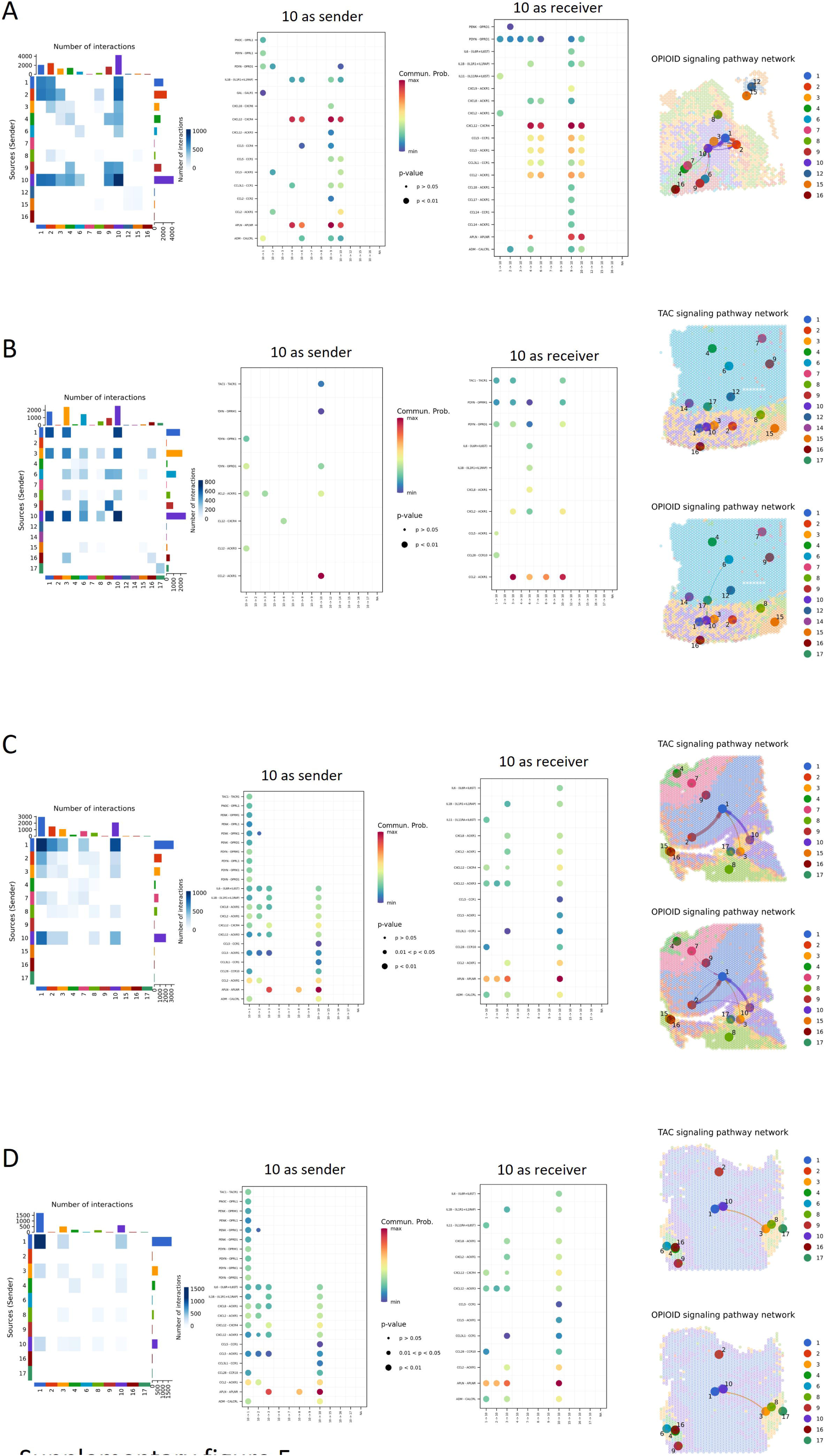
For each sample: left panel shows heatmap created using CellChatV2 showing number of R-L interactions between each cluster; middle panels show bubble plots created using CellChatV2 showing R-L communication probability for selected R-L pathways in NH18-1756, where left is for cluster 10 is the sender (ligand is expressed by cluster 10) and right is for cluster 10 as receiver (receptor is in cluster 10); right panel shows **s**patial plots for TAC signalling and opioid signalling created using CellChatV2 showing R-L interactions between each cluster. Thickness of arrows represents number of interactions. RT14 is shown in **A**; RT13 is shown in **B**; RT6 is shown in **C**; RT1 is shown in **D**.

**Supplementary figure 6:**
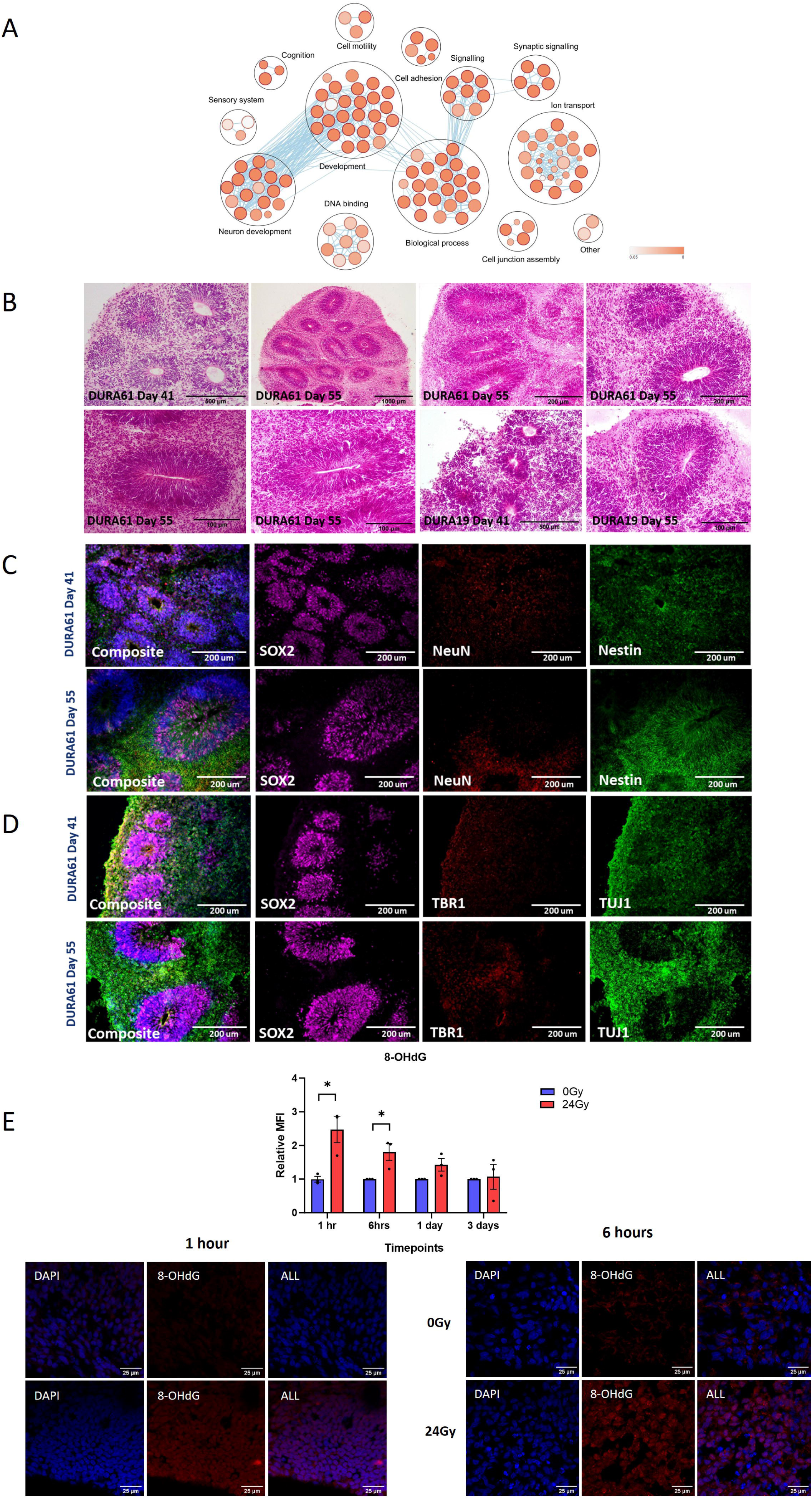
**A** GO analysis using all (844) DMGs shared between patient samples and COs. Edge cutoff 0.6. **B** H&E staining of COs from day 41 and day 55 of maturation from two patient-derive EPSC lines (DURA61 and DURA19). **C** Representative IF images of COs at day 41 and 55 stained with antibodies against SOX2, NeuN, Nestin and DAPI. Scale bars are 200 μm. **D** Representative IF images of COs at day 41 and 55 stained with antibodies against SOX2, TBR1, TUJ1 and DAPI. Scale bars are 200 μm. **E** Top panel show bar plot depicting relative mean fluorescence intensity (MFI) of 8-OHdG comparing irradiated organoids (red) to the control (blue) at timepoints ranging from 1 hour to 3 days post-irradiation. Each data point represents pooled data from a single CO. Unpaired t-test was performed to assess statistical significance. Error bars represent SEM. Bottom panels show representative IF images of 8-OHdG (red) at 1 hour (left) and 6 hours (right) post-irradiation. Nuclei were counterstained for DAPI (blue). Scale bas are 25 μm.

**Supplementary figure 7:**
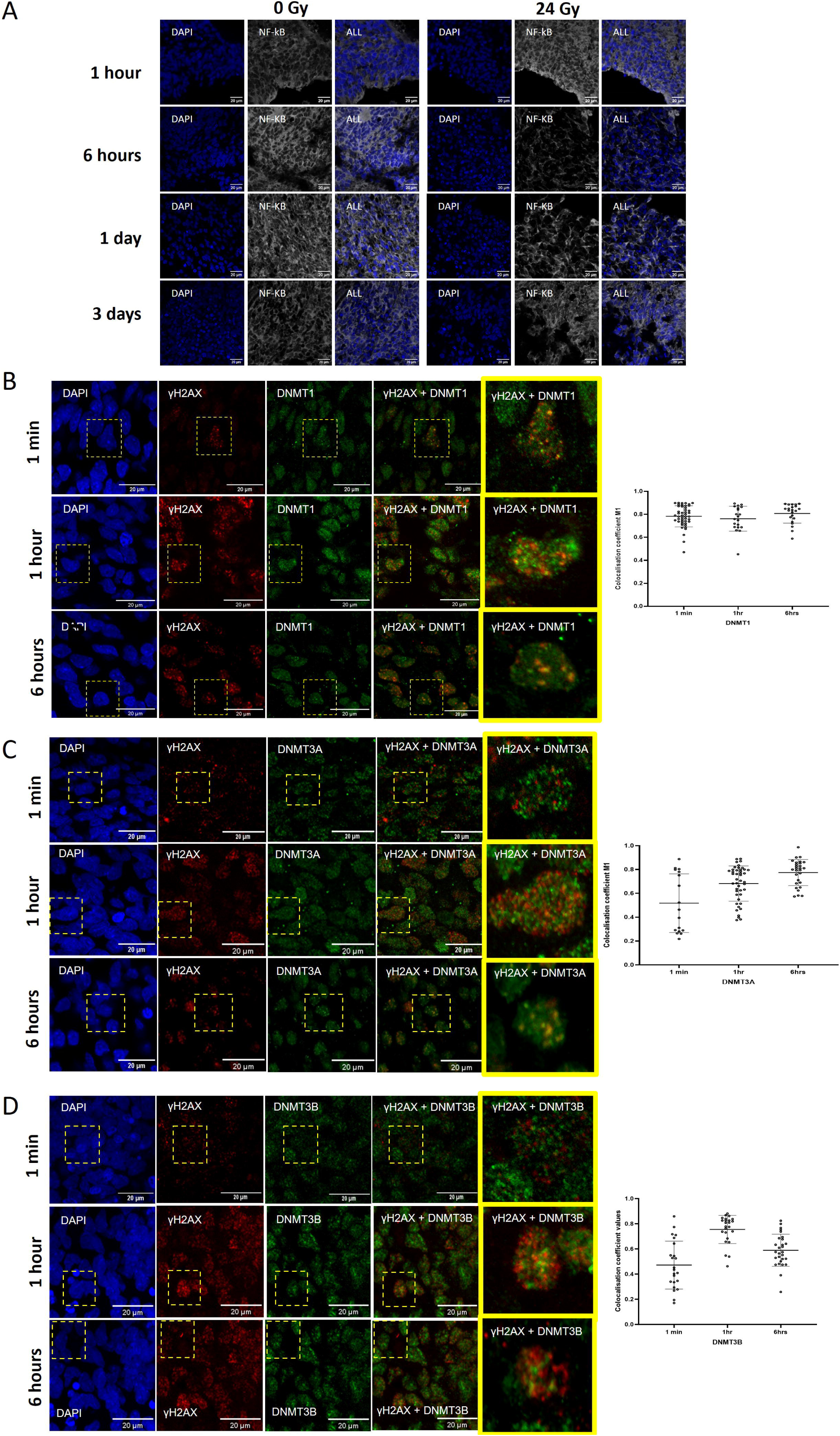
**A** Representative IF images of NF-kB p65 (white) at 1 hour, 6 hours, 1 day and 3 days post-irradiation. Nuclei were counterstained for DAPI (blue). Scale bas are 20 μm. **B-D** Left panels show representative immunofluorescence (IF) images illustrating colocalisation of γH2AX (red) with DNMT1 (**A**), DNMT3A (**B**) and DNMT3B (**C**) (all green) at 1 minute, 1 hour and 6 hours post-irradiation. The overlay images demonstrate points of colocalisation in yellow. Nuclei were counterstained for DAPI (blue). Scale bars are 20um. Right panels show dot plots of Manders colocalisation coefficient (M1) value from individual nuclei displayed from organoids at each timepoint. The error bars represent SD.

**Supplementary figure 8:**
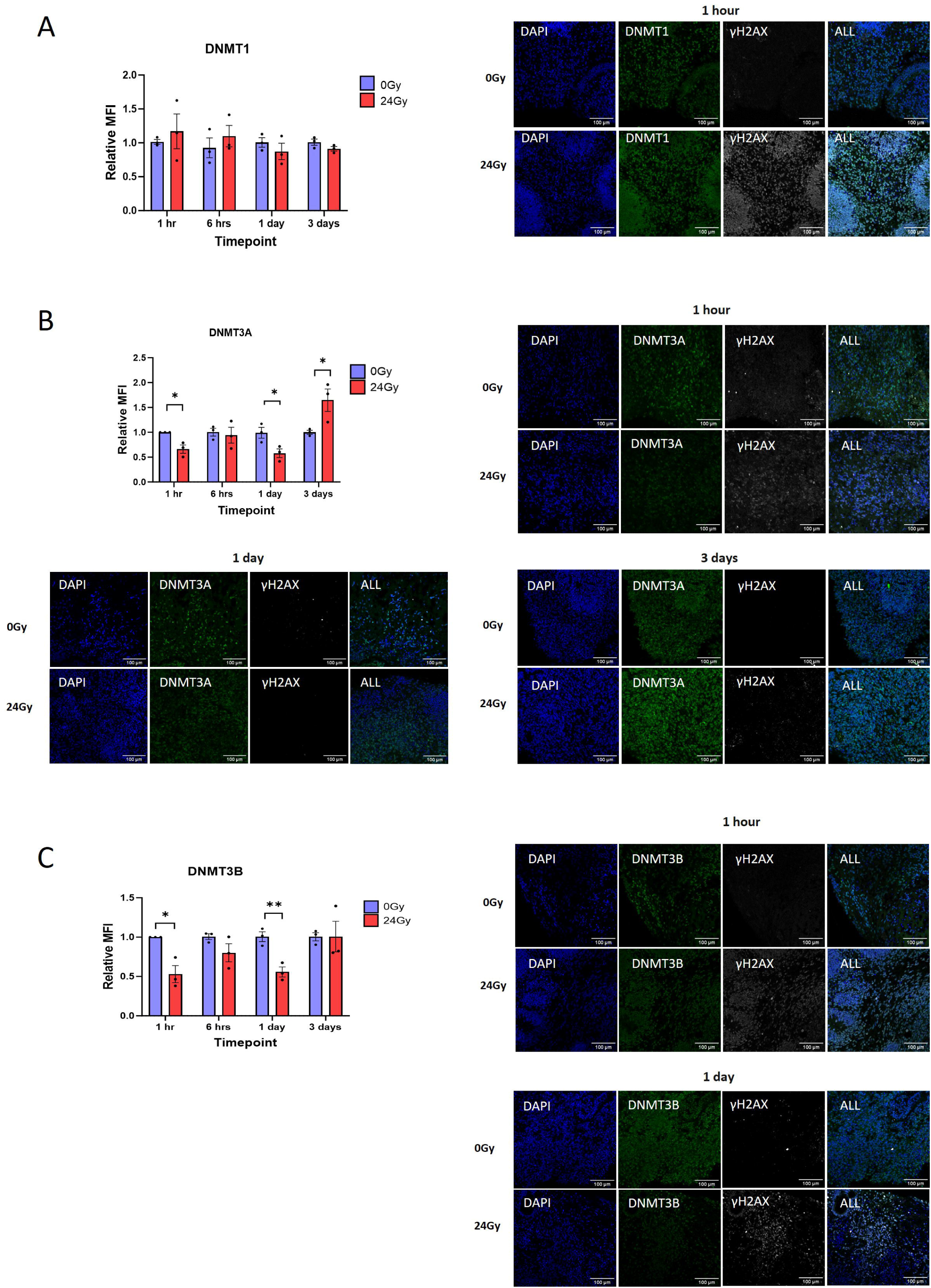
Top left panels are those seen in figure 6J, showing bar plots depicting relative mean fluorescence in tensity (MFI) of DMNT1 (**A**), DNMT3A (**B**) and DNMT3B (**C**) at timepoints ranging from 1 hour to 3 days post-irradiation. Each data point on the graph represents pooled data from a single CO. Unpaired t-test was performed to assess statistical significance. Error bars represent the standard error of the mean (SEM). Other panels depict representative IF images of DMNT1 (**A**), DNMT3A (**B**) and DNMT3B (**C**) (all green) and γH2AX (white) at various timepoints post-irradiation.

